# Genetic interaction mapping highlights key roles of the Tol-Pal complex

**DOI:** 10.1101/2021.09.13.460050

**Authors:** Wee Boon Tan, Shu-Sin Chng

## Abstract

The conserved Tol-Pal trans-envelope complex is important for outer membrane (OM) stability and cell division in Gram-negative bacteria. It has been proposed to mediate OM constriction during cell division via tethering to the cell wall. Yet, recent studies suggest that the complex has additional roles in OM lipid homeostasis and septal cell wall separation. How the Tol-Pal complex functions to facilitate these many processes is unclear. To gain insights into its role(s), we applied transposon insertion sequencing, and report here a detailed network of genetic interactions with the *tol-pal* locus in *Escherichia coli*. We found one positive and >20 negative strong interactions based on fitness. Disruption of genes responsible for osmoregulated periplasmic glucan biosynthesis restores fitness and OM barrier function, but not cell division defects, in *tol-pal* mutants. In contrast, deletions of genes involved in OM homeostasis and cell wall remodelling give rise to synthetic growth defects in strains lacking Tol-Pal, especially exacerbating OM barrier and/or cell division defects. Notably, the Δ*tolA* mutant having additional defects in OM protein assembly (Δ*bamB*) exhibited severe division phenotypes, even under conditions where the single mutants divide normally; this highlights the possibility for OM phenotypes to indirectly influence the cell division process. Overall, our work provides insights into the intricate nature of Tol-Pal function, and reinforces the model that this complex plays crucial roles in cell wall-OM tethering, cell wall remodelling, and in particular, OM homeostasis.

## Introduction

The complex Gram-negative bacterial cell envelope comprises the inner membrane (IM), the cell wall, and the outer membrane (OM) [1]. For proper growth and division, the synthesis and expansion of all three layers have to be tightly regulated and coordinated. In particular, the biosynthesis and maintenance of the cell wall and OM, being physically separated from the cytoplasm, require more specialized mechanisms. The cell wall or peptidoglycan polymer is synthesized in the periplasm by SEDS (shape, elongation, division and sporulation) proteins and PBPs (penicillin binding proteins), and is cleaved by hydrolases to facilitate remodeling during cell growth and division [2]. To build the OM, dedicated protein systems including the Lpt bridge, the Bam machine, and the Lol pathway are required to transport and assemble lipopolysaccharides (LPS), OM β-barrel proteins, and OM lipoproteins, respectively [3-5]. How bulk phospholipids (PLs) are trafficked between the membranes is less well understood [6]. In *Escherichia coli*, the cell wall and OM are also tethered covalently by lipoprotein Lpp, and non-covalently by lipoprotein Pal and β-barrel protein OmpA, for stability and function [7]. It remains unclear how cell wall and OM assembly and homeostasis are coordinated during growth and division.

The Tol-Pal complex is a conserved trans-envelope system that appears to be involved in both cell wall and OM homeostasis [8-10]. It consists of an IM complex containing TolQ, TolR, and TolA and an OM complex composed of periplasmic protein TolB and Pal. The TolQRA complex is energized by the IM proton motive force (pmf), and interacts with TolB, and possibly Pal, via the periplasmic domain of TolA [11-14]. Cells lacking the Tol-Pal complex exhibit pleiotropic phenotypes [12, 14]. Specifically, these strains exhibit hypersensitivity towards antibiotics and leak periplasmic contents, indicative of a defective OM. The same cells also bleb OM vesicles primarily from the septum, pointing at a possible cell division defect. Consistently, *tol-pal* mutants are known to form undivided chains, but only under osmotic stress, and are particularly sensitive to such stress at high temperatures [10, 15]. The complex nature of these phenotypes has made it difficult to assign Tol-Pal function. Nevertheless, it has been shown that the Tol proteins localize to the septum during cell division, and are required for Pal to accumulate at the cell division site – this enrichment of septal cell wall-OM tethering is believed to be important for coordinated OM invagination, and may account for the OM hypervesiculation phenotype [15-17]. Recently, Tol-Pal was also suggested to modulate cell wall remodeling; chained mutant cells at low osmolarity have unseparated peptidoglycan sacculus, and overexpressing certain cell wall hydrolases can suppress this phenotype [10]. Even so, OM defects are still present in these suppressors, as is the case when cells are not osmotically stressed. Importantly, it has been demonstrated that mutants lacking an intact Tol-Pal complex accumulate excess PLs at the OM and display slower retrograde PL transport, potentially accounting for the origin of the unstable/defective OM under all growth conditions [8, 18]. Taken together, the Tol-Pal complex is seemingly involved in a plethora of diverse processes: peptidoglycan-OM tethering, cell wall remodeling, and retrograde PL transport.

The function(s) of the Tol-Pal complex remains elusive. It is believed that TolQRA dynamically controls TolB-Pal interaction to facilitate Pal binding to septal cell wall [9], but how the complex affects cell wall remodeling and PL trafficking is not clear. One possibility is that peptidoglycan-OM tethering is the main function of the Tol-Pal complex, loss of which somehow gives rise to problems in the latter two processes. However, it is known that deletion of *pal* yields milder phenotypes relative to the deletion of other *tol* genes [10], implying that the function(s) of Tol-Pal might not be centered around the ability of Pal to bind peptidoglycan but instead depends largely on TolQRA and/or TolB. The Tol proteins may possibly rely on additional interacting partner(s), or be somewhat sufficient, to facilitate cell wall remodeling and PL trafficking processes. It is conceivable that these processes are independent functions of the Tol complex. It is also possible that disruption of one process indirectly results in the other. Notably, preventing cell chaining of *tol* mutants does not alleviate OM permeability defects [10]; therefore, OM instability, likely due to defects in retrograde PL trafficking, is not a downstream effect of problems in cell wall remodeling. The molecular function(s) of the Tol-Pal complex, and the relationships between the existing phenotypes, remain to be clarified.

In this study, we adopted an unbiased genetic approach to gain additional insights into the function(s) of the Tol-Pal complex. Using high-throughput transposon insertion sequencing [19], we mapped and validated positive and negative genetic interactions, based on fitness, with the *tol*-*pal* locus in *E. coli*. We found only one strong positive interaction; abolishing biosynthesis of osmo-regulated periplasmic glucans (OPGs) improves fitness and OM barrier function in the Δ*tolA* mutant, but does not rescue its cell division defects. In contrast, we identified >20 negative genetic interactions with *tolQ* and *tolA*. These genes are primarily involved in cell wall and OM processes, consistent with the fact that the Tol-Pal complex plays crucial roles in cell wall remodeling and OM lipid homeostasis. Interestingly, we noted that additional OM defects that are not known to impact cell division can exacerbate the division defects in Δ*tolA* strains, suggesting it is possible for OM phenotypes to indirectly affect cell division. Notwithstanding the complex relationships between various *tol-pal* effects in cell wall-OM tethering, cell wall remodeling and lipid homeostasis, our work reinforces the idea that the Tol-Pal complex may play a primary role in OM structure and function.

## Results

### *En masse* transposon directed insertion sequencing (TraDIS) reveals extensive genetic interactions with *tol*-*pal*

Transposon insertion site mapping, using methods such as Tn-seq or TraDIS, have been widely applied to study essentiality and fitness impact of genes in different organisms [20, 21]. This technique has also been extended to investigate synthetic interactions between genes via comparative analysis of transposon insertion site profiles in different strain backgrounds [22, 23]. To clarify its physiological role(s), we applied TraDIS to identify all possible genetic interactions with the *tol-pal* locus. Any gene with increased or decreased frequency of transposon insertion in the *tol*-*pal* but not the WT background would indicate positive or negative genetic interaction with the locus, respectively (Fig. 1A). We generated separate Tn5 transposon mutant libraries in *E. coli* WT, Δ*tolQ*, and Δ*tolA* strains. Each library contains approximately one million Tn5 mutants, as estimated from the total numbers of colonies obtained from transposition. Three replicate cultures were grown from each library in LB media, and genomic DNA were isolated and prepared for sequencing using an optimized TraDIS protocol adapted from Langridge et al. [24]. From a single multiplexed sequencing run (MiSeq) of the nine samples, a total of ∼6.7 million Tn5 junction-containing reads were recovered, with ∼1.9, ∼2.9, and ∼1.7 million combined reads from the WT, Δ*tolQ*, and Δ*tolA* triplicates, respectively (Fig. 1B; Table S1). Alignment of sequences adjacent to the insertion junctions allowed pinpointing of the exact insertion sites on the genome. We identified at least 430,000 unique insertion sites in each background, with one insertion every 9-11 bp on average, indicating that our Tn5 libraries have extensive coverage of all possible non-essential genes (Figs. 1B, S1A).

**Figure 1.**
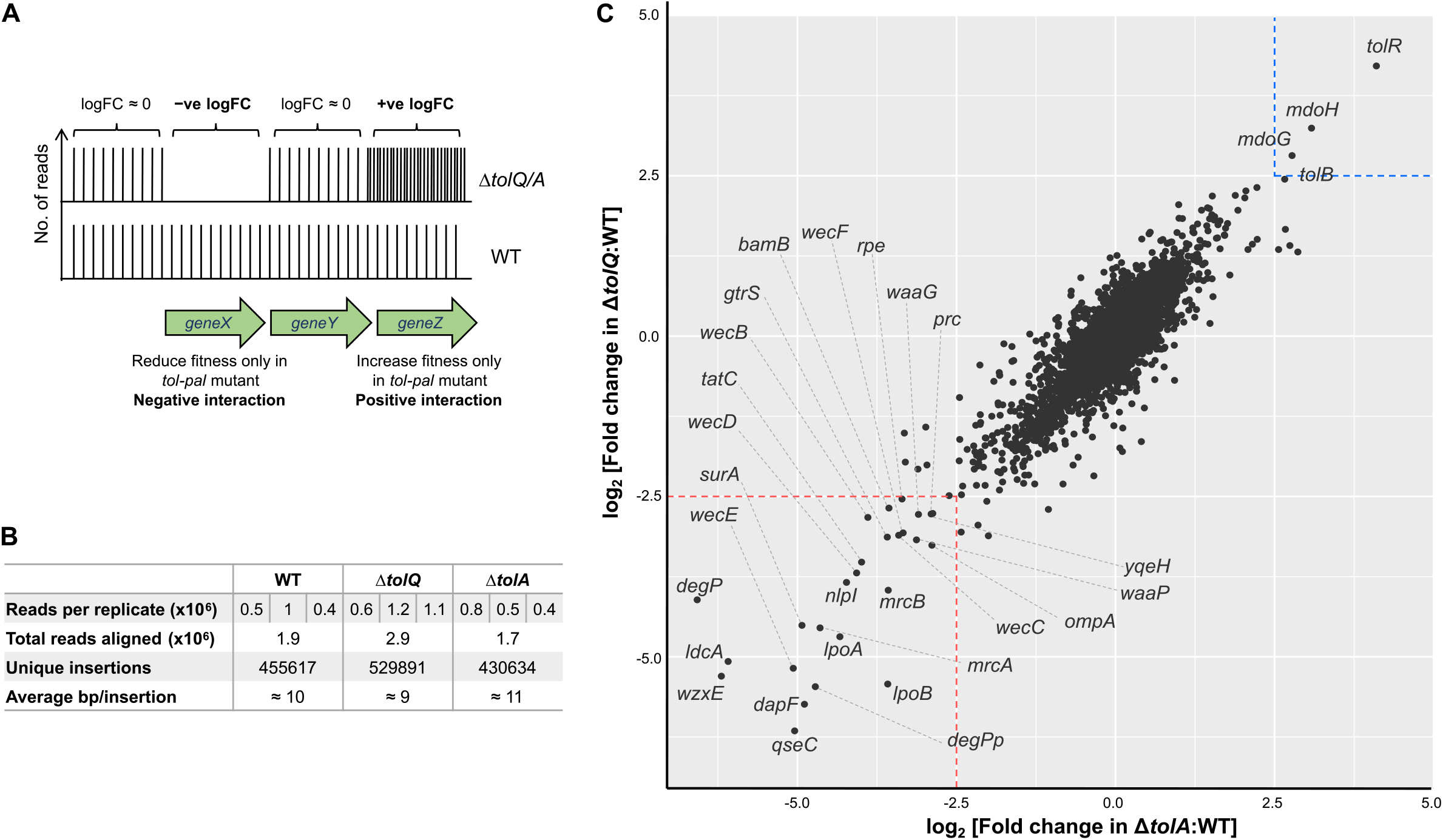
Saturated genetic interaction map with the *tol-pal* locus. (A) Schematic representation of the rationale of using transposon insertion site sequencing (TraDIS) to identify genetic interactions. Increased or decreased frequency of insertion within a gene in Δ*tolQ/A* background relative to WT represents positive or negative genetic interactions, respectively. (B) Summary of the total number of sequencing reads recovered that aligned to the MC4100 reference genome [53]. Numbers of reads for each replicate, total combined reads, and number of unique insertion sites in the genome for each background are indicated. (C) Comparison of all genetic interactions specific to Δ*tolQ* and Δ*tolA* backgrounds. Genes that exhibit strong positive or negative genetic interaction with both *tolQ* and *tolA* (arbitrary threshold of log_2_FC>2.5 or <-2.5, corrected P-value < 10^−5^) are annotated.

To identify genes that exhibited strong interactions, we compared the total number of reads for all insertions in each gene in the Δ*tolQ*/*A* background to that in WT. Genes that contained less than ten insertion reads in both WT and mutant backgrounds were removed from analysis. In either the Δ*tolQ*:WT or Δ*tolA*:WT comparison, a number of genes displayed statistically significant (FDR corrected P-value <10^−5^) log_2_ fold change (log_2_FC) value of >2.5 in either direction (Figs. S1B, C; Tables S2, S3), indicative of strong genetic interactions with *tolQ* or *tolA*. In fact, the comparative profiles were well correlated (Fig. 1C; R^*2*^ = 0.76 for 3912 genes), suggesting that most genetic interactions are common, and therefore specific to overall Tol-Pal function(s). We noted that *tolR/A/B* and *tolQ/R/B* were among the genes with strongest enrichment of transposon insertion reads in Δ*tolQ* and Δ*tolA* backgrounds, respectively, relative to WT (Figs. S1B-D). This is expected given that these genes are from the same pathway, thus disruption would only affect WT but no longer have additional deleterious effects in Δ*tolQ*/*A* backgrounds. Intriguingly, insertion frequency in *pal* remained unchanged in either Δ*tolQ* or Δ*tolA* relative to WT (log_2_FC of -0.08 or +0.3, respectively; statistically insignificant, Figs. S1B-D), perhaps suggesting less major contribution to overall function of the complex. Using the stringent cutoff (|log_2_FC| > 2.5), our TraDIS screen uncovered 2 other genes showing strong positive genetic interactions with both *tolQ* and *tolA*, and 25 genes exhibiting common negative interactions.

### Disrupting biosynthesis of osmoregulated periplasmic glucans results in positive genetic interactions with *tol-pal*

Transposon insertions in *mdoH* (*opgH*) and *mdoG* (*opgG*), two genes necessary for the biosynthesis of osmoregulated periplasmic glucans (OPGs) [25], were strongly enriched in both Δ*tolQ* and Δ*tolA* backgrounds (Figs. 1C, 2A), suggesting improved fitness. A third gene involved in but not required for OPG biosynthesis, *mdoD* (*opgD*), also exhibited positive interactions with *tolQ/A*, albeit with lower fold-enrichment in insertion frequency (Table S4). Interestingly, we have previously found that mutations in *mdoH* may contribute to suppression of vancomycin sensitivity in *tol-pal* mutants [26]. We proceeded to validate this interaction by examining both growth and vancomycin sensitivity of a Δ*tolA* strain with *mdoH* deleted. The Δ*tolA* Δ*mdoH* double mutant indeed grew better (as judged by optical density measurements), and had restored vancomycin resistance compared to the Δ*tolA* strain (Figs. 2B, C). Mutants lacking the Tol-Pal complex also exhibit cell division defects; however, Δ*mdoH* did not rescue the cell chaining nor the plating defects (at 37°C) observed in the Δ*tolA* strain under low osmolarity conditions (Figs. 2C, D). MdoH has a moonlighting function in growth rate regulation via sequestration of FtsZ by its N-terminal domain [27]; we showed that expression of the N-terminal domain *in trans* did not reverse the suppression (Fig. S2), suggesting that the positive genetic interaction observed was unrelated to FtsZ sequestration. We conclude that removing OPGs from the periplasm somehow confers better growth and OM barrier function in mutants lacking Tol-Pal.

**Figure 2.**
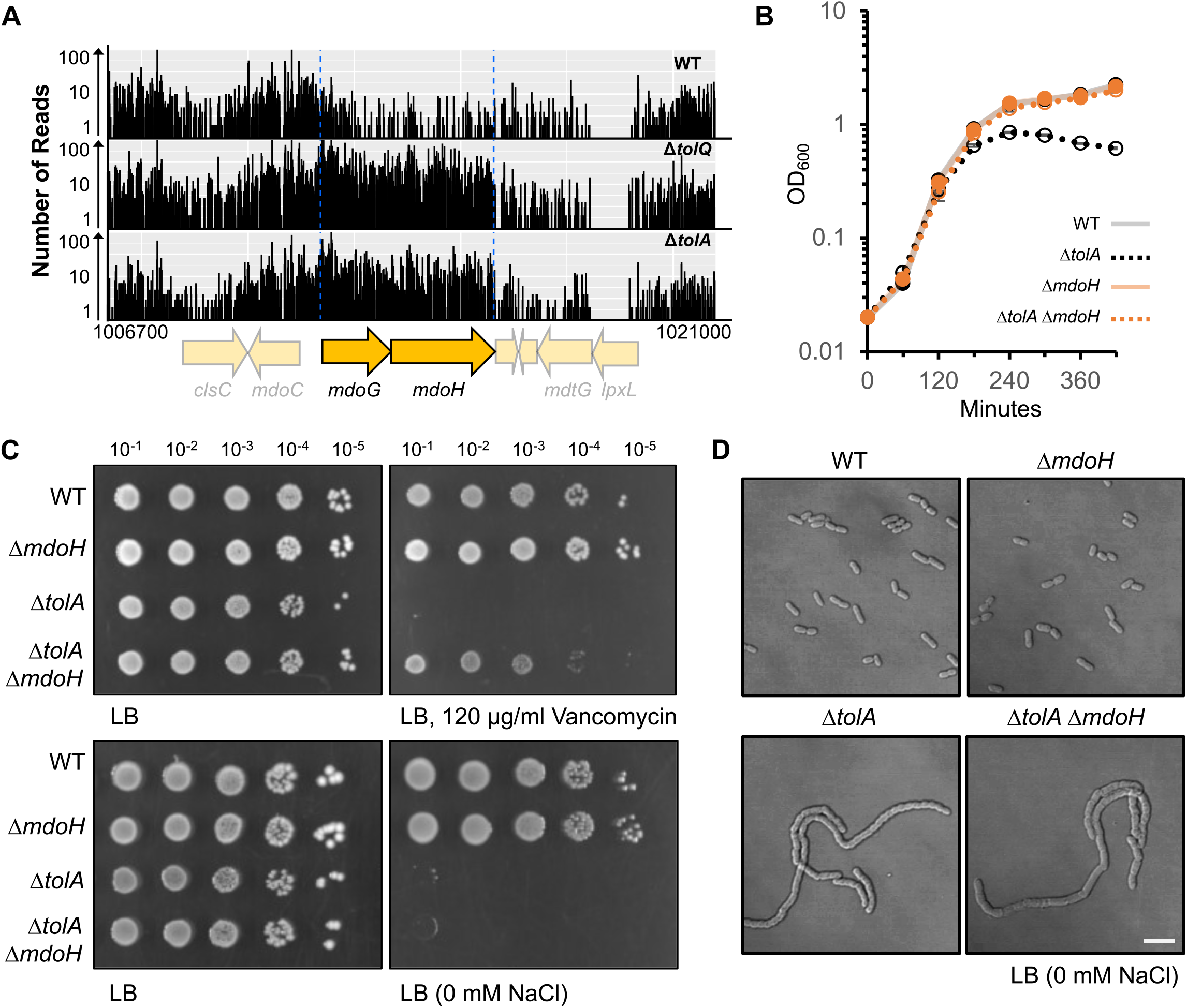
Δ*mdoH* restores fitness and OM barrier function but not cell division defects in the Δ*tolA* strain. (A) Transposon insertion profiles at the *mdoGH* locus in WT, Δ*tolQ* and Δ*tolA* backgrounds. Numbers of reads (raw data before normalization for downstream analysis) at indicated positions on the genome are plotted with open reading frames annotated below the horizontal axis. (B) Growth curves of indicated strains, based on optical density at 600 nm, in LB media at 37°C. (C) Efficiency of plating (EOP) of indicated strains on LB media without NaCl, or containing vancomycin, at 37°C. (D) Differential interference contrast (DIC) images of cells from indicated strains grown to mid-log phase in LB media without NaCl at 37°C. Scale bar represents 5 µm.

### Genes exhibiting negative interactions with *tol-pal* are important for outer membrane or cell wall integrity

25 genes contained significantly fewer insertions in both Δ*tolQ* and Δ*tolA* background relative to WT, indicative of negative genetic interactions (Fig. 1C). Among these are several genes that are known to exhibit synthetic phenotypes with *tol-pal* (*bamB, degP, lpoA, lpoB, mrcA (PBP1A), mrcB (PBP1B), ompA, surA*) [28-30], validating our overall TraDIS approach. Generally, Δ*tolQ* and Δ*tolA* mutants appeared less tolerant of disruption in genes involved in two major processes, namely cell wall biosynthesis/remodeling and OM assembly/integrity (Table 1). For cell wall biosynthesis/remodeling, we specifically found genes involved in peptidoglycan synthesis (*dapF, lpoA, mrcA, lpoB, mrcB*), remodeling (*nlpI, prc, tatC*), recycling (*ldcA*), and undecaprenyl lipid carrier sequestration (*qseC, wecB, wecC, wecD, wecE, wecF, wzxE* [31, 32]); for OM assembly/integrity, genes involved in LPS synthesis/modification (*gtrS, waaP, waaG*), β-barrel protein assembly (*bamB, degP, surA*), and OM-cell wall tethering (*ompA*) exhibited negative interactions with *tol-pal*. We successfully constructed clean deletions of each gene in the Δ*tolA* background, with the exception of *qseC*, where a depletion strain was constructed (Fig. S3). We demonstrated that most of these double mutants displayed synthetic growth defects when grown in LB liquid media at 37°C (Figs. S3, S4) and many exhibited reduced plating efficiencies on M9 minimal (37°C) or LB (30°C, 37°C, 42°C) media (Figs. S3, S5). Based on the growth phenotypes, the more severe negative interactions were further characterized below.

**Table 1.** Negative genetic interactions with the *tol-pal* locus.

|  | Category | Genes |
| --- | --- | --- |
| Cell wall<br>biosynthesis or<br>remodelling | Peptidoglycan biosynthesis | *dapF, lpoA, mrcA, lpoB, mrcB* |
|  | Cell wall remodelling | *nlpl, prc, tatC* |
|  | Cell wall recycling | *ldcA* |
|  | Und-P lipid carrier related | *qseC, wecB, wecC, wecD, wecE, wecF, wzxE* |
| OM assembly or<br>integrity | LPS synthesis/modification | *gtrS, waaG, waaP* |
|  | β-barrel protein assembly | *bamB, degP, surA* |
|  | OM-cell wall tethering | *ompA* |
| Other |  | *rpe, yqeH* |

### Mutants lacking Tol-Pal are sensitive to defects in cell wall remodeling

Among the genes important for cell wall integrity, *nlpI, prc*, and *tatC* exhibited the most severe negative interactions with *tol-pal* (Figs. S4, S5, 3A-C). The OM lipoprotein NlpI physically regulates the tail-specific protease Prc to primarily degrade MepS, a peptidoglycan endopeptidase important for cell wall expansion [33]. Disruption of *nlpI* or *prc* in the Δ*tolA* background resulted in severe growth defects in liquid cultures (Fig. 4A). We noticed that the overnight stationary phase cultures used as inoculums had 10-to 100-fold fewer colony-forming unit (CFU) per OD, indicating that the double mutants lost viability during late stationary phase (Fig. 4B, C). Consistent with this idea, microscopic images of these cells revealed extensive amounts of cell debris, as well as morphological defects, only at late stationary phase (Fig. 5), corresponding to when the proteolytic effect of NlpI/Prc on MepS is most crucial [33]. Incidentally, we observed slight increases in the frequency of Tn5 insertion in *mepS* in the *ΔtolQ* and Δ*tolA* backgrounds relative to WT (log_2_FC of +1.16 or +1.07, respectively; Tables S2, S3), suggesting that MepS activity could be unfavorable in strains lacking Tol-Pal. The synthetic defects with Δ*tolA* observed here were likely due to problems related to elevated MepS activity in the absence of NlpI and Prc.

TatC is part of the twin-arginine translocation (Tat) system important for the secretion of folded-proteins containing Tat-specific signal peptides [34, 35]. Deletion of *tatC* is known to result in cell chaining and envelope defects, largely due to impaired translocation of cell wall amidases AmiA and AmiC [34]. Due to the chaining phenotype, the Δ*tatC* single mutant had ∼10-fold fewer CFU per OD compared to WT, despite similar growth profiles (Figs. 4D, E). The Δ*tolA* Δ*tatC* double mutant grew drastically slower than either single mutant in LB media, and formed much longer unseparated chains, consistent with a strong negative genetic interaction (Fig. 5). The severe cell division defects likely account for why the double mutant was unable to grow on low osmolarity media at 30°C, a condition where the single mutants can survive (Fig. 4F). We also observed that the Δ*tolA* Δ*tatC* strain exhibited hypersensitivity towards vancomycin (Fig. 4F). TatC functions with TatB, TatA and/or TatE to transport substrates; these three proteins are structurally homologous and have partially overlapping roles [35, 36], especially TatA and TatE, possibly explaining why we did not observe negative interactions between these genes with *tolQ*/*A*. Given that mutants lacking the Tol-Pal complex already exhibit problems in OM invagination and/or septal cell wall separation [10, 15], it makes sense that removing TatC further exacerbates defects in cell wall remodeling during cell division, thus giving rise to the synthetic interaction.

### *tol-pal* mutants cannot withstand additional OM perturbation

Mutants lacking functional Tol-Pal complex are known to exhibit severe OM defects likely due to lipid dyshomeostasis [8]. It appears that further disruption of the OM, particularly defects in OMP assembly, LPS synthesis, and OM-cell wall tethering, are not well-tolerated in Δ*tolQ* and Δ*tolA* strains (Fig. 3D-G). The BAM machine comprises essential β-barrel protein BamA and lipoprotein BamD, and three non-essential lipoproteins BamB, BamC and BamE [3]. We only observed strong negative interactions between *tolQ*/*A* and *bamB*, but not *bamC*/*E*, likely because removing either of the latter genes has relatively weaker impact on OMP assembly [37, 38]. Removing BamB resulted in severe growth defects in the Δ*tolA* background in both liquid culture and solid media (Figs. 6A, C); similar synthetic sick phenotypes were observed in Δ*pal* Δ*bamB* mutant previously [30]. Despite increasing optical density, CFUs recoverable from the liquid culture of the Δ*tolA* Δ*bamB* mutant were extremely low and did not increase over time (Fig. 6B). While the cellular morphology of Δ*tolA* and Δ*bamB* single mutants are largely similar to that of WT in LB media, we found that the Δ*tolA* Δ*bamB* double mutant exhibited severe chaining phenotype (Fig 7), which may explain the disconnect between CFU and OD. This observation is somewhat surprising given that defects in OMP assembly are not known to impact cell division.

**Figure 3.**
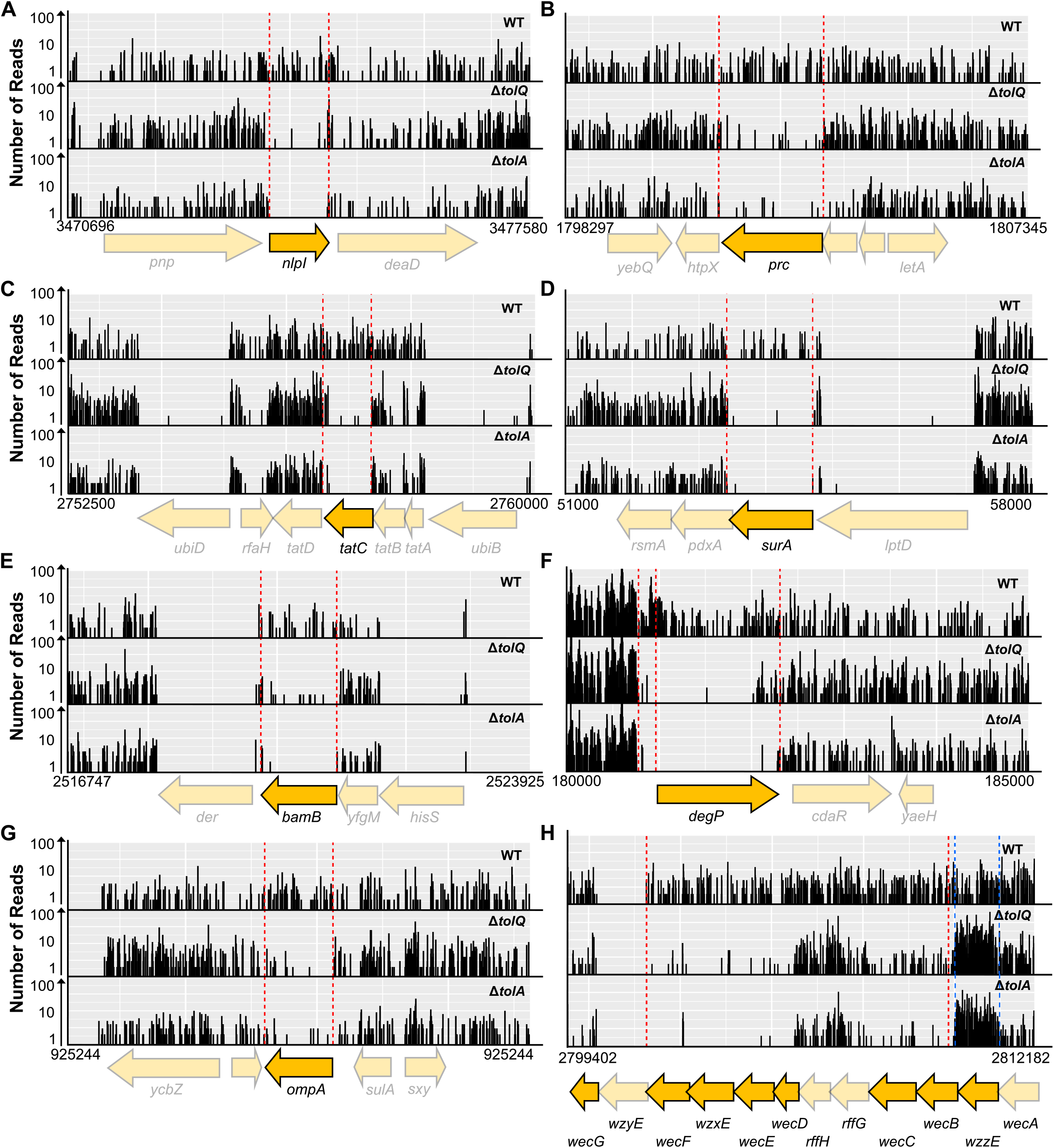
The frequencies of transposon insertions decrease significantly in genes involved in cell wall and OM homeostasis in *tol-pal* strains. Transposon insertion profiles at selected loci that exhibited strong negative interactions with *tolQ* and *tolA*, including (A) *nlpI*, (B) *prc*, (C) *tatC*, (D) *surA*, (E) *bamB*, (F) *degP*, (G) *ompA*, and (H) ECA biosynthesis gene cluster.

In addition to *bamB*, both *surA* and *degP*, which encode for the main OMP periplasmic chaperone and protease [3], respectively, also exhibited negative genetic interactions with *tol-pal* (Fig. 1C). We showed that the Δ*tolA* Δ*surA* double mutant displayed a growth defect in liquid LB media at 37°C, and a severe plating defect on solid media at 30°C (Figs. 6A, C). Removing SurA did not further sensitize the Δ*tolA* strain towards SDS EDTA but did increase vancomycin sensitivity (Fig. S6A). Interestingly, the Δ*tolA* Δ*surA* double mutant was somehow more vancomycin-resistant than the Δ*surA* mutant alone. The Δ*tolA* Δ*degP* double mutant also displayed a growth defect in liquid LB media at 37°C (Fig. 6A); however, as previously reported, this strain could not grow on plate at 37°C (Fig. 6C; [28]). At the permissive temperature of 30°C, the Δ*tolA* Δ*degP* double mutant was more sensitive to SDS EDTA, but not vancomycin, when compared to the Δ*tolA* strain (Fig. S6B). Actively growing Δ*tolA* Δ*surA* and Δ*tolA* Δ*degP* cells also exhibited chaining phenotypes, albeit milder than observed in the Δ*tolA* Δ*bamB* strain (Fig. 7). Taken together, while the synthetic interactions uncovered are necessarily complex, our data indicate a clear requirement for proper OMP biogenesis and homeostasis in strains lacking Tol-Pal.

Defects in the LPS biosynthesis pathway also appear to exhibit synthetic phenotypes with *tol-pal*. Specifically, disruption of *waaG* and *waaP* resulted in slightly slower growth and exacerbated OM permeability defects, particularly vancomycin sensitivity, in the Δ*tolA* strain (Fig. S7A). The Δ*tolA* Δ*waaG* mutant especially exhibited cell morphological changes in logarithmic and stationary growth phases (Figs. S7B, C). WaaG is responsible for the installation of the first glucose in the outer core of LPS [39]; Δ*waaG* mutants have truncated LPS structure, and thus a deep rough phenotype. WaaP is a kinase that phosphorylates the first heptose residue in the LPS, absence of which affects other phosphorylation and glycosylation events in the inner core [39]. Other genes involved in LPS core biosynthesis (*waaB, waaO, waaQ*) also had reduced Tn5 insertions in both Δ*tolQ* and Δ*tolA* background, although not meeting our initial threshold of log_2_FC<-2.5 (Table S2, S3). Therefore, the OM of *tol-pal* mutants also cannot tolerate additional perturbations to LPS core structure that may affect LPS packing in the outer leaflet.

Loss of OmpA, the abundant OMP that non-covalently tethers OM to peptidoglycan [40], caused one of the worst growth defects in mutants lacking Tol-Pal (Figs. 1, S4). The Δ*tolA* Δ*ompA* mutant grew much slower in liquid culture (Fig. 6D) and displayed lowered plating efficiency across all growth phases (Fig. 6E); these phenotypes are likely due to the gross morphological defects observed in this strain (Fig. 7). Interestingly, removing Lpp, which links the OM and cell wall covalently [7], has no impact on growth of Δ*tolQ*/*A* strains in our TraDIS experiment (Table S2, S3), suggesting that the observed synthetic interactions were not due to general loss of OM-peptidoglycan tethering. Deletion of *ompA* causes general OM defects in the forms of reduced stiffness and increased sensitivity to antibiotics [41, 42]. We infer that *tol-pal* mutants could not tolerate disturbance to OM stability due to specific loss of OmpA (see discussion).

## Discussion

The specific function(s) of the Tol-Pal complex has been difficult to assign [8, 10, 15]. Here, using TraDIS, we have mapped the complex genetic interaction network with the *tol-pal* locus (Fig. 1). We showed that in *tol-pal* mutants, disrupting biosynthesis of OPGs (Δ*mdoG/H*) resulted in the only positive interaction (based on stringent cutoff of log_2_FC>2.5), where fitness and OM barrier, but not division defects, were largely restored (Fig. 2). In contrast, we observed multiple negative interactions with *tol-pal* almost exclusively due to further defects in either the cell wall or OM (Table 1; Figs 4-7); Δ*tolA* strains exhibited strong synthetic defects especially when combined with mutations in cell wall remodelling (Δ*nlpI*, Δ*prc*, and Δ*tatC*) or OM assembly (Δ*bamB*, Δ*degP*, Δ*surA*, Δ*waaG*, and Δ*waaP*) pathways. Overall, our results are consistent with the idea that the Tol-Pal complex plays important roles in OM homeostasis and cell wall remodelling, in addition to OM-peptidoglycan tethering.

**Figure 4.**
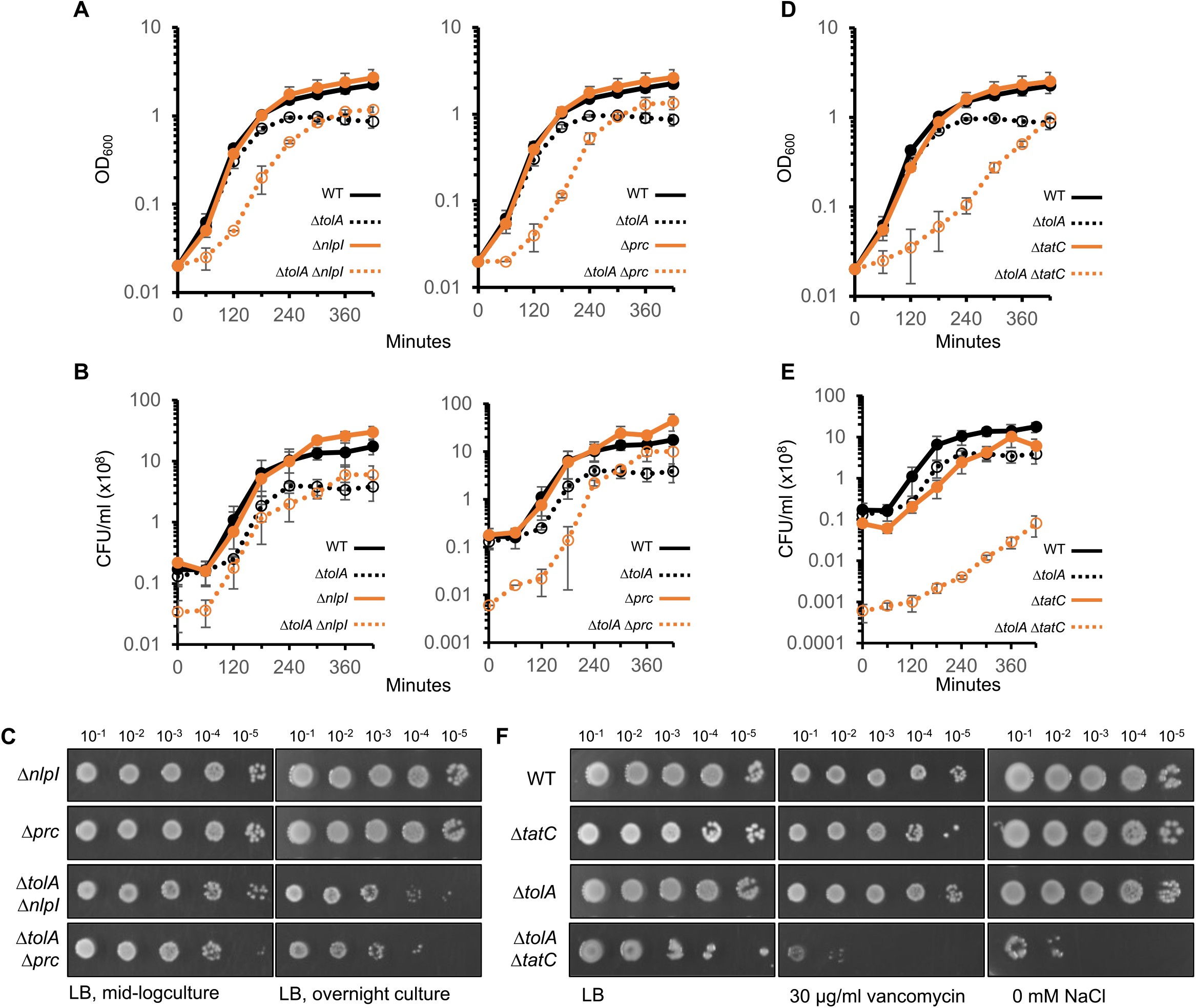
Cell wall remodeling defects are detrimental to cells lacking TolA. (A) Growth curves of indicated *nlpI*/*prc* strains, based on optical density at 600 nm, in LB media at 37°C. (B) Colony-forming-unit (CFU) enumeration of indicated *nlpI*/*prc* strains at the corresponding timepoints taken during growth curve in (A). (C) EOP of indicated *nlpI*/*prc* strains from either mid-log or overnight cultures on LB media at 37°C. Cultures were normalized to the same initial optical density. (D) Growth curves of indicated *tatC* strains, based on optical density at 600 nm, in LB media at 37°C. (E) CFU enumeration of indicated *tatC* strains at the corresponding timepoints taken during growth curve in (D). (F) EOP of indicated *tatC* strains on LB media without NaCl, or containing vancomycin, at 37°C.

**Figure 5.**
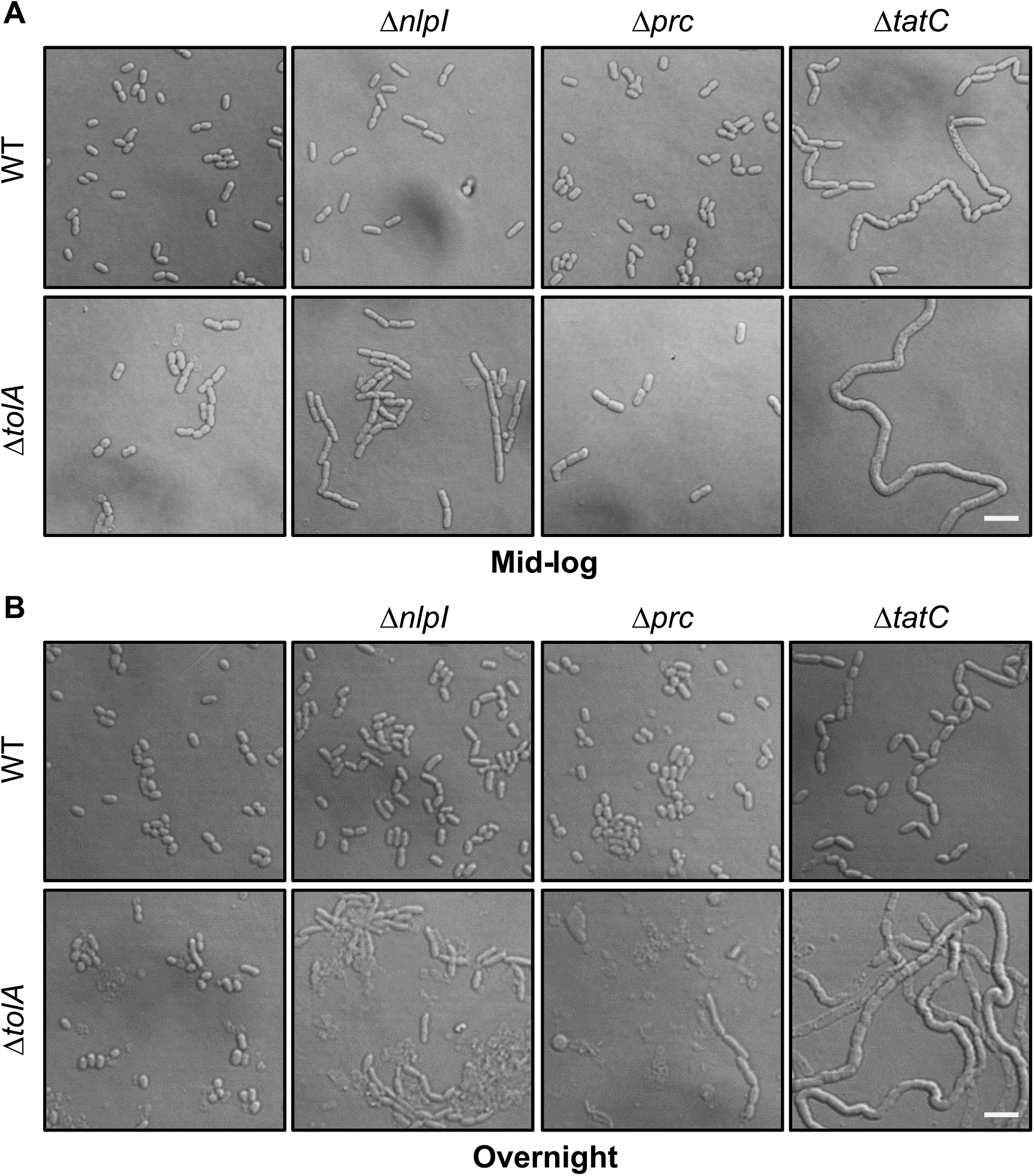
Δ*tolA* Δ*nlpI*, Δ*tolA* Δ*prc*, and Δ*tolA* Δ*tatC* double mutants exhibit morphological defects. DIC images of cells from indicated strains grown to (A) mid-log or (B) stationary phase in LB media (with NaCl) at 37°C. Scale bar represents 5 µm.

**Figure 6.**
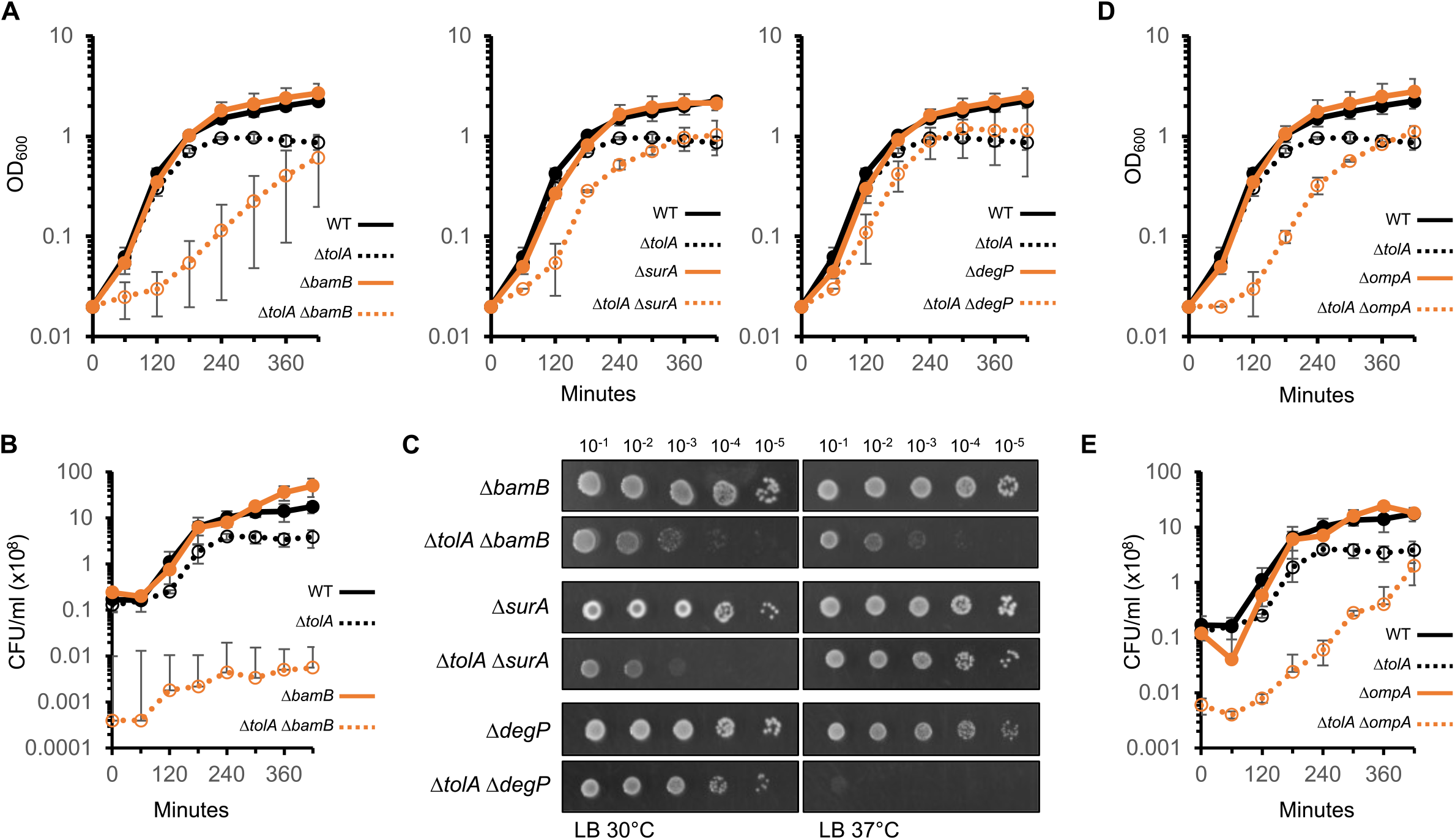
Defects in OM biogenesis and stability are detrimental in Δ*tolA* strains. (A) Growth curves of indicated *bamB*/*surA*/*degP* strains, based on optical density at 600 nm, in LB media at 37°C. (B) CFU enumeration of indicated *bamB* strains at the corresponding timepoints taken during growth curve in (A). (C) EOP of indicated *bamB*/*surA*/*degP* strains on LB media at either 30°C or 37°C. (D) Growth curves of indicated *ompA* strains, based on optical density at 600 nm, in LB media at 37°C. (E) CFU enumeration of indicated *ompA* strains at the corresponding timepoints taken during growth curve in (D).

**Figure 7.**
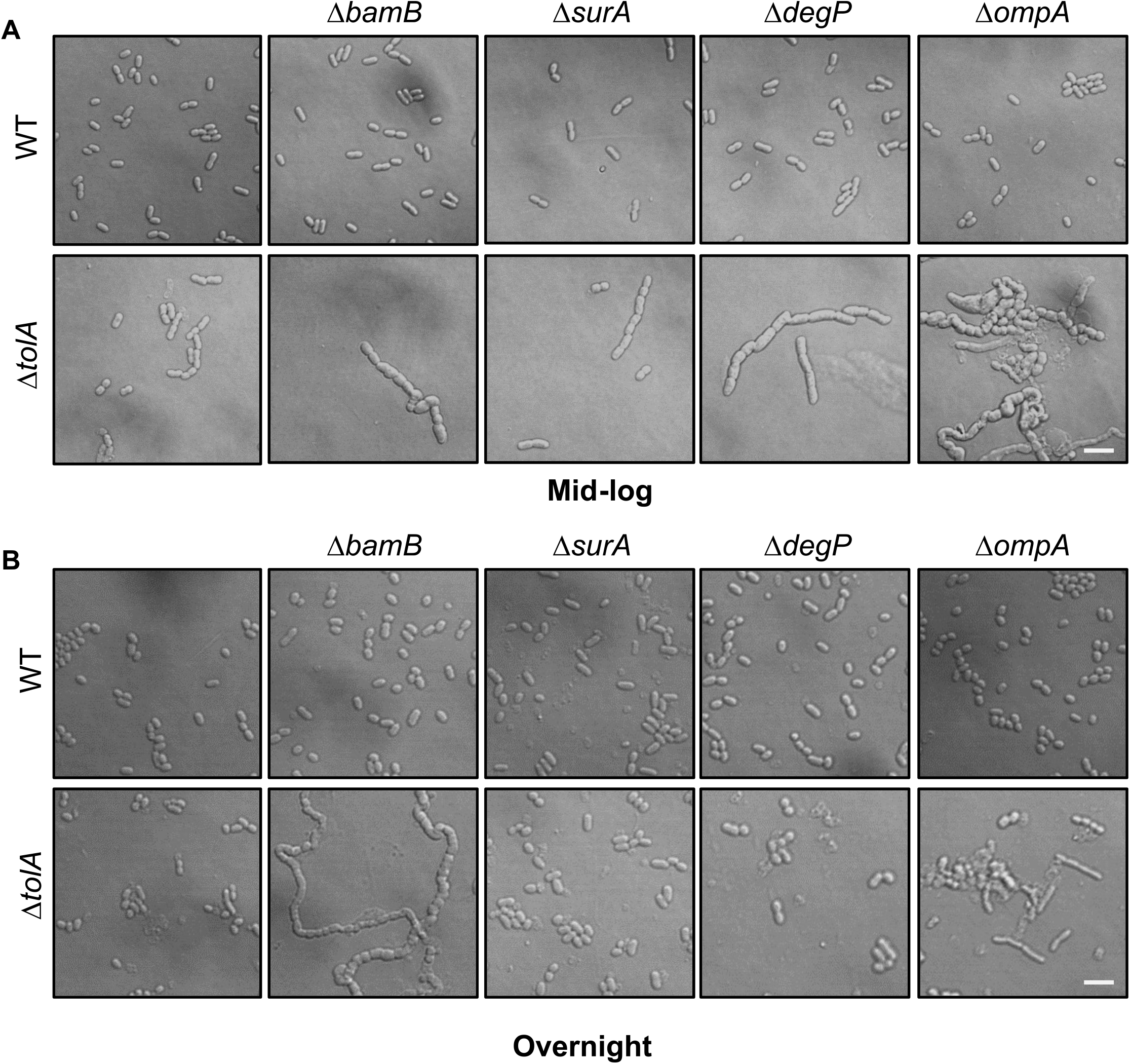
Δ*tolA* Δ*bamB*, Δ*tolA* Δ*surA*, Δ*tolA* Δ*degP*, and Δ*tolA* Δ*ompA* double mutants exhibit morphological defects. DIC images of cells from indicated strains grown to (A) mid-log or (B) stationary phase in LB media (with NaCl) at 37°C. Scale bar represents 5 µm.

An interesting observation in our TraDIS dataset is that transposon insertions in *pal* were not enriched in the Δ*tolQ/A* backgrounds relative to WT (Figs. S1B-D). Intuitively, deleting any additional gene in the *tol-pal* locus should be neutral in Δ*tolQ/A* mutants but has a strong detrimental effect in the WT strain. This is true for the other *tol* genes, as judged by the increase in insertion frequencies in these genes in the Δ*tolQ/A* strains over WT, but clearly not the case for *pal*. Therefore, it appears that deletion of *pal* does not impact the fitness of the WT strain in the same way as removing other *tol* genes. This is consistent with the milder phenotypes Δ*pal* strains have displayed [10]. Consequently, septal peptidoglycan-OM tethering offered by Pal may be peripheral to the main function of the TolQRA-TolB system.

Besides the *tol* genes, it is intriguing that *mdoG*/*H* were the only other genes that exhibited strong positive interactions with Δ*tolQ*/*A*. Removing MdoH restored fitness and OM barrier function of mutants lacking Tol-Pal, but how the resulting loss of OPGs suppresses these phenotypes remains elusive. Nevertheless, it is known that removal of OPGs improves OM barrier function in general [26, 43, 44], suggesting that the suppression effect might not be specific to the *tol-pal* locus, and therefore less related to its direct function. At a less stringent cutoff (log_2_FC > 1.25), additional positive genetic interactions with both Δ*tolQ* and Δ*tolA* are revealed (Table S4). While we did not go on to validate these TraDIS interactions, we note that higher insertion frequencies were observed in genes involved in cell division (*minC, minD, minE*), lipid homeostasis (*cdh, fadR, kdsC*), cell wall homeostasis (*ddlB, elyC, ldtB*) and others (e.g., *wzzE, yhdP*). Additional characterization of these putative interactions would be warranted to understand their connections to the Tol-Pal complex.

In an earlier study, we have demonstrated that accumulation of intermediates in the enterobacterial common antigen (ECA) pathway is linked to restoration of OM barrier function in strains lacking Tol-Pal [26]. The fact that Tn5 insertions within *wzzE* were enriched in Δ*tolQ* and Δ*tolA* backgrounds (log_2_FC of +1.68 and +1.73, respectively; Fig. 3H) is consistent with our observation that deletion of *wzzE* restores OM barrier function in the Δ*tolA* strain. However, several ECA biosynthesis genes that restored OM barrier function when deleted (*wecB, wecC, wecE, wecF*) were found to exhibit strong negative interactions with *tolQ*/*A* (log_2_FC < -2.5). This is not unexpected given that the build-up of ECA-related intermediates is known to result in cell wall defects via the sequestration of undecaprenyl-phosphate (und-P) [32], the lipid carrier also used in peptidoglycan biosynthesis; indeed, *wec tol* double mutants do display more severe morphological defects than the corresponding single mutants [26]. One of the strongest negative genetic interaction with *tolQ* and *tolA* is *qseC*, encoding the sensor kinase of the QseBC two-component system, that has recently been discovered to be important for cell wall homeostasis, especially when und-P pool is limited [31]. Interestingly, the same genetic screen for mutants sensitive to a limited und-P pool also identified *wec* and *tol-pal* genes, overall suggesting that *tol-pal* strains are sensitive to cell wall defects linked to und-P sequestration.

The observed negative interaction between Δ*ompA* and Δ*tolQ/A* is particularly strong. Since TolQRA/B contributes to localization of Pal to the septal cell wall, this genetic interaction suggests some level of redundancy between Pal and OmpA, which contain homologous peptidoglycan binding domains [40]. Consistently, synthetic interactions between Δ*pal* and Δ*ompA* has previously been reported [30]. However, it is not clear how OmpA can contribute during cell division. While TolB has been shown to bind OmpA [45], the latter is not freely diffusible in the OM to facilitate recruitment to the cell septum [46]. An alternative reasoning is that cells cannot tolerate the simultaneous loss of OM-cell wall tethering at the cell septum and around the cell, respectively. Yet, in our Δ*tolA* Δ*ompA* strains, Pal and Lpp are still present, and therefore should still be able to contribute to OM-cell wall tethering in the cell periphery in place of OmpA. We also did not observe any synthetic interactions between *tolQ/A* and *lpp*; the requirement of OmpA in *tol* strains is therefore puzzling. We posit that the loss of OmpA gives rise to specific OM defects that are not well tolerated in strains with existing OM lipid dyshomeostasis.

Clearly, *tolQ/A* mutants could not tolerate deletion of genes that affect peptidoglycan biosynthesis and remodelling, or OM structure and assembly. The simplest explanation for the interactions with cell wall processes is that mutants lacking functional Tol-Pal already have cell septum separation defects during cell division, albeit only under conditions of low osmolarity. Likewise, the synthetic phenotypes with OM defective mutants are possibly due to existing OM lipid dyshomeostasis in *tol-pal* strains. Notably, one of the strongest interactions we have observed was with OMP assembly, specifically Δ*bamB*. The severe growth defects detected in the double Δ*tolA* Δ*bamB* mutant are likely due to a combination of OM lipid imbalance (Δ*tolA*), and OMP assembly defects (Δ*bamB*). Like Δ*tolA*, Δ*bamB* mutant also contains excess PLs in the OM [8], indicating the double mutant has exacerbated OM lipid dyshomeostasis. One interesting phenotype in the double mutant is the severe chaining defect in cells grown under conditions of normal osmolarity, where either Δ*tolA* or Δ*bamB* alone do not chain. These results overall indicate that exaggerated OM (lipid) problems can give rise to issues during cell division. While our data does not rule out the possibility that cell wall separation and OM lipid homeostasis may be two largely independent functions of Tol-Pal, it appears to be consistent with the idea that maintaining proper OM lipid balance could be the primary function, thus indirectly affecting septa separation. Although the mechanism(s) by which Tol-Pal (or specifically TolQRA and/or TolB) facilitate these functions remains enigmatic, our work reinforces the emerging roles of Tol-Pal in ensuring proper cell wall separation during cell division and, importantly, in maintaining proper OM lipid homeostasis in Gram-negative bacteria.

## Experimental procedures

### Strains, plasmid, and growth conditions

All the strains used in this study are listed in Table S5. *Escherichia coli* strain MC4100 [*F-araD139* Δ*(argF-lac) U169 rpsL150 relA1 flbB5301 ptsF25 deoC1 ptsF25 thi*] [47] was used as the wild-type (WT) strain for most of the experiments. NR754, an *araD*^+^ revertant of MC4100 [48], was used as the WT strain for experiments involving arabinose-inducible vector. pBAD43-*tolA* was cloned using the EcoRI and XbaI restriction sites with primers EcoRI-*tolA5* (ACAATGAATTCCGAGAGTGTCAAAGGCAACC) and XbaI-*tolA3* (TACTGTCTAGATTACGGTTTGAAGTCCAATGGC). pTrc99a-*mdoH*-N pTrc99a-*mdoH*-FL were cloned using NcoI and XbaI restriction sites with the 5’ primer NcoI-*mdoH5* (TTTTCCATGGCAAATAAGACAACTGAGTACATT) in combination with either the XbaI-*mdoH*-N (TTTTTCTAGATTAGTGATGGTGATGGTGATGTCCGCCACCACGGCGGATGGTACCGACG) or XbaI-*mdoH*-FL (TTTTTCTAGATTAGTGATGGTGATGGTGATGTCCGCCACCTTGCGAAGCCGCATCCGG), respectively. Gene deletion mutants were constructed using recombineering [49, 50]. Primers used are listed in Table S6. Whenever needed, the antibiotic resistance cassettes were flipped out as described [50]. Gene deletion cassettes were transduced into relevant genetic background strains via P1 transduction [51]. Strains were grown in Luria-Bertani (LB) broth (1% tryptone and 0.5% yeast extract, with or without 1% NaCl). Agar plates contain 1.5% agar in the corresponding media.

### Transposon mutant library construction

Recipient strains of *E. coli* were grown to mid-log, washed 4 times with cold sterile H_2_O, and concentrated 100x with cold sterile H_2_O to generate electrocompetent cells. For each transposon mutagenesis reaction, 100µl of competent cells was mixed with 1 µl of EZ-Tn5 KAN-2 Tnp Transposome (Epicentre) and electroporated at 1800V. 1ml SOC media was added immediately after electroporation and the mixture was allowed for recovery at 37°C for 2 hours. The recovered reaction was plated onto multiple LB plates (depending on the efficiency of each strain) with 50 µg/ml kanamycin and incubated overnight at 37°C. On subsequent day, number of colonies per reaction was estimated and the colonies were scrapped and pooled in LB with 15% glycerol and stored at -80°C. Final mutant libraries (estimated ≈ 1,000,000 mutants) were generated by pooling multiple libraries of the same background.

### Multiplexed TraDIS

Genomic DNAs were isolated from LB culture inoculated with the respective library after 3 hours growth at 37°C. For each mutant library, three cultures were grown separately for genomic DNA isolation (triplicate). For each sample, 100 ng of genomic DNA was sheared with Covaris and then subsequently end-repaired and -adenylated following Illumina TruSeq Nano protocol. Ligation of sequencing adapters with indices specific to each sample was also done according to TruSeq Nano kit protocol. Each adapter-ligated library was enriched for Tn5 containing fragment with 2-steps PCR using KAPA Library Amplification Kit at 65°C reannealing temperature for 10 (1^st^ amplification) or 20 (2^nd^ amplification) cycles and 500nM final concentration for each primer. First round of PCR was done with a 24 bases forward primer that contains the exact sequence of the -47 to -24 (relative to the junction of transposon insertion) region of the Tn5 transposon and a reverse primer that is complimentary to the 3’ region of the TruSeq indexed adapter. Second round of PCR was done with the same reverse primer and an 86 bases forward primer that contains, from 5’ to 3’, the 58 bases TruSeq universal adapter sequence, a 6 bases additional index sequence (different for each sample), and a 22 bases sequence that contains the exact sequence of the -32 to -11 region of the Tn5 transposon. Primers used for TraDIS sample preparation are listed in Table S7. Nine libraries were pooled and normalized to 10nM and 20% PhiX spike-in was added. The pooled library was loaded onto MiSeq at 8pM final concentration and sequenced using 300 cycles, 2×150 paired end protocol.

### TraDIS analysis

The ESSENTIALS pipeline [52] was used to analyze the output of TraDIS sequencing. Demultiplexed R1 reads from MiSeq sequencing were uploaded to the ESSENTIALS server and analyzed using the following setting: Tn insertion = random; library size = 1,000,000; filtering non-unique sequence = yes; barcode mismatch allowed = 0; genomic sequence length after removal of barcode and transposon = 100; barcode = beginning of line; transposon inverted repeat = beginning of line; minimal sequence match for alignment = 50; strand to align = both; use truncated version of gene = yes; remove genomic position bias using Loess = yes; normalization method = TMM; analysis = qCML; modelling of variance = tagwise; prior.n = 5; p-value adjustment = BH. Through this analysis, all reads were filtered for the correct barcode and Tn5 sequence, and 100 bases right after the Tn5 insertion junction were aligned to an annotated MC4100 reference genome (NZ_HG738867.1) [53]. Each mapped read indicates a Tn5 insertion at the exact position of the genome. The frequency of Tn5 insertion in each gene were normalized and compared between Δ*tolQ* or Δ*tolA* library against the WT control library. Genes with less than 10 reads mapped in all three replicates of both mutant and WT library were removed from the analysis and were not compared between Δ*tolQ* and Δ*tolA* backgrounds. Downstream data was inspected in Microsoft Excel and graphical visualizations of the data were plotted using the *ggplot2* package in R.

### Growth curve and CFU enumeration

To evaluate growth, overnight cultures were first normalized to OD_600_ = 0.02 into fresh LB media. For measurement with plate reader, 200 µl of the diluted cultures were transferred onto 96-well plate and OD_490_ at 37 °C was measured at fixed interval using Victor X4 plate reader (Perkin Elmer). For manual growth curve and CFU enumeration, at 1-hour intervals, OD_600_ for the culture at 37°C were measured and a small volume of the culture were serially diluted and plated to estimate the CFU/ml at that time point.

### Efficiency of plating assay

Efficiency of plating (EOP) analyses were done by spotting serially diluted culture onto different agar plates. Cells cultures were 10-fold serially diluted in 150mM NaCl on 96-well plate. Two or five microliters of each dilution were spotted onto the plates and incubated overnight at the indicated temperatures. All results shown are representative of at least three independent replicates.

### Microscopy

For each strain, five microliters of overnight or mid-log cultures were spotted onto freshly prepared 1% agarose pads made with the corresponding media. Images were acquired using a Zeiss LSM710 confocal microscope at 100x magnification.

## Acknowledgements

We thank Suk Yean Poon and William Burkholder (Institute of Molecular Cell Biology, A*STAR) for their generous help with MiSeq sequencing. We are grateful to Varnica Khetrapal, Kurosh Mehershahi, Shannon Fenlon, and Swaine Chen (Genome Institute of Singapore, A*STAR) for guidance in sample preparation and initial analyses. We also thank Swaine Chen and Lok-To Sham (National University of Singapore) for critical comments on the manuscript. W.B.T. was supported by the NUS Integrative Science and Engineering Program-SCELSE scholarship. This work was supported by Singapore Ministry of Health National Medical Research Council under its Open Fund Individual Research Grant (MOH-000145) (to S.-S.C.). The authors declare no conflict of interest.

## References

1. Nikaido, H., Molecular basis of bacterial outer membrane permeability revisited. Microbiology and molecular biology reviews, 2003. 67(4): p. 593–656.

2. Egan, A.J., J. Errington, and W. Vollmer, Regulation of peptidoglycan synthesis and remodelling. Nature Reviews Microbiology, 2020. 18(8): p. 446–460.

3. Hagan, C.L., T.J. Silhavy, and D. Kahne, β-Barrel membrane protein assembly by the Bam complex. Annual review of biochemistry, 2011. 80: p. 189–210.

4. Okuda, S., et al., Lipopolysaccharide transport and assembly at the outer membrane: the PEZ model. Nature Reviews Microbiology, 2016. 14(6): p. 337–345.

5. Okuda, S. and H. Tokuda, Lipoprotein sorting in bacteria. Annual review of microbiology, 2011. 65: p. 239–259.

6. Shrivastava, R. and S.-S. Chng, Lipid trafficking across the Gram-negative cell envelope. Journal of Biological Chemistry, 2019. 294(39): p. 14175–14184.

7. Asmar, A.T. and J.-F. Collet, Lpp, the Braun lipoprotein, turns 50—major achievements and remaining issues. FEMS microbiology letters, 2018. 365(18): p. fny199.

8. Shrivastava, R., X.E. Jiang, and S.S. Chng, Outer membrane lipid homeostasis via retrograde phospholipid transport in Escherichia coli. Molecular microbiology, 2017. 106(3): p. 395–408.

9. Szczepaniak, J., C. Press, and C. Kleanthous, The multifarious roles of Tol-Pal in Gramnegative bacteria. FEMS Microbiology Reviews, 2020. 44(4): p. 490–506.

10. Yakhnina, A.A. and T.G. Bernhardt, The Tol-Pal system is required for peptidoglycan-cleaving enzymes to complete bacterial cell division. Proceedings of the National Academy of Sciences, 2020. 117(12): p. 6777–6783.

11. Cascales, E., et al., Proton motive force drives the interaction of the inner membrane TolA and outer membrane pal proteins in Escherichia coli. Molecular microbiology, 2000. 38(4): p. 904–915.

12. Lloubès, R., et al., The Tol-Pal proteins of the Escherichia coli cell envelope: an energized system required for outer membrane integrity? Research in microbiology, 2001. 152(6): p. 523–529.

13. Sturgis, J.N., Organisation and evolution of the tol-pal gene cluster. Journal of molecular microbiology and biotechnology, 2001. 3(1): p. 113–122.

14. Bernadac, A., et al., Escherichia coli tol-pal mutants form outer membrane vesicles. Journal of bacteriology, 1998. 180(18): p. 4872–4878.

15. Gerding, M.A., et al., The trans-envelope Tol–Pal complex is part of the cell division machinery and required for proper outer-membrane invagination during cell constriction in E. coli. Molecular microbiology, 2007. 63(4): p. 1008–1025.

16. Petiti, M., et al., Tol energy-driven localization of Pal and anchoring to the peptidoglycan promote outer-membrane constriction. Journal of molecular biology, 2019. 431(17): p. 3275–3288.

17. Szczepaniak, J., et al., The lipoprotein Pal stabilises the bacterial outer membrane during constriction by a mobilisation-and-capture mechanism. Nature communications, 2020. 11(1): p. 1–14.

18. Masilamani, R., M.B. Cian, and Z.D. Dalebroux, Salmonella Tol-Pal reduces outer membrane glycerophospholipid levels for envelope homeostasis and survival during bacteremia. Infection and immunity, 2018. 86(7): p. e00173–18.

19. Barquist, L., et al., The TraDIS toolkit: sequencing and analysis for dense transposon mutant libraries. Bioinformatics, 2016. 32(7): p. 1109–1111.

20. Goodall, E.C., et al., The essential genome of Escherichia coli K-12. MBio, 2018. 9(1): p. e02096–17.

21. Rifat, D., et al., Genome-Wide Essentiality Analysis of Mycobacterium abscessus by Saturated Transposon Mutagenesis and Deep Sequencing. Mbio, 2021. 12(3): p. e01049–21.

22. Truong, T.T., A. Vettiger, and T.G. Bernhardt, Cell division is antagonized by the activity of peptidoglycan endopeptidases that promote cell elongation. Molecular Microbiology, 2020. 114(6): p. 966–978.

23. Winkle, M., et al., DpaA detaches Braun’s lipoprotein from peptidoglycan. Mbio, 2021. 12(3): p. e00836–21.

24. Langridge, G.C., et al., Simultaneous assay of every Salmonella Typhi gene using one million transposon mutants. Genome research, 2009. 19(12): p. 2308–2316.

25. Bontemps-Gallo, S., J.-P. Bohin, and J.-M. Lacroix, Osmoregulated periplasmic glucans. EcoSal Plus, 2017. 7(2).

26. Jiang, X.E., et al., Mutations in enterobacterial common antigen biosynthesis restore outer membrane barrier function in Escherichia coli tol-pal mutants. Molecular Microbiology, 2020. 114(6): p. 991–1005.

27. Hill, N.S., et al., A moonlighting enzyme links Escherichia coli cell size with central metabolism. PLoS genetics, 2013. 9(7): p. e1003663.

28. Kern, B., O.P. Leiser, and R. Misra, Suppressor mutations in degS overcome the acute temperature-sensitive phenotype of Δ degP and Δ degP Δ tol-pal mutants of Escherichia coli. Journal of bacteriology, 2019. 201(11): p. e00742–18.

29. Typas, A., et al., Regulation of peptidoglycan synthesis by outer-membrane proteins. Cell, 2010. 143(7): p. 1097–1109.

30. Typas, A., et al., High-throughput, quantitative analyses of genetic interactions in E. coli. Nature methods, 2008. 5(9): p. 781–787.

31. Jorgenson, M.A. and J.C. Bryant, A genetic screen to identify factors affected by undecaprenyl phosphate recycling uncovers novel connections to morphogenesis in Escherichia coli. Molecular Microbiology, 2021. 115(2): p. 191–207.

32. Jorgenson, M.A., et al., Dead-end intermediates in the enterobacterial common antigen pathway induce morphological defects in Escherichia coli by competing for undecaprenyl phosphate. Molecular microbiology, 2016. 100(1): p. 1–14.

33. Singh, S.K., et al., Regulated proteolysis of a cross-link–specific peptidoglycan hydrolase contributes to bacterial morphogenesis. Proceedings of the National Academy of Sciences, 2015. 112(35): p. 10956–10961.

34. Ize, B., et al., Role of the Escherichia coli Tat pathway in outer membrane integrity. Molecular microbiology, 2003. 48(5): p. 1183–1193.

35. Palmer, T. and P.J. Stansfeld, Targeting of proteins to the twin-arginine translocation pathway. Molecular microbiology, 2020. 113(5): p. 861–871.

36. Ize, B., et al., In vivo dissection of the Tat translocation pathway in Escherichia coli. Journal of molecular biology, 2002. 317(3): p. 327–335.

37. Sklar, J.G., et al., Lipoprotein SmpA is a component of the YaeT complex that assembles outer membrane proteins in Escherichia coli. Proceedings of the National Academy of Sciences, 2007. 104(15): p. 6400–6405.

38. Wu, T., et al., Identification of a multicomponent complex required for outer membrane biogenesis in Escherichia coli. Cell, 2005. 121(2): p. 235–245.

39. Bertani, B. and N. Ruiz, Function and biogenesis of lipopolysaccharides. EcoSal Plus, 2018. 8(1).

40. Park, J.S., et al., Mechanism of anchoring of OmpA protein to the cell wall peptidoglycan of the gram-negative bacterial outer membrane. The FASEB Journal, 2012. 26(1): p. 219–228.

41. Rojas, E.R., et al., The outer membrane is an essential load-bearing element in Gram-negative bacteria. Nature, 2018. 559(7715): p. 617–621.

42. Smani, Y., et al., Role of OmpA in the multidrug resistance phenotype of Acinetobacter baumannii. Antimicrobial agents and chemotherapy, 2014. 58(3): p. 1806–1808.

43. Murakami, K., et al., The Absence of Osmoregulated Periplasmic Glucan Confers Antimicrobial Resistance and Increases Virulence in Escherichia coli. Journal of bacteriology, 2021. 203(12): p. e00515–20.

44. Stokes, J.M., et al., Cold stress makes Escherichia coli susceptible to glycopeptide antibiotics by altering outer membrane integrity. Cell chemical biology, 2016. 23(2): p. 267–277.

45. Clavel, T., et al., TolB protein of Escherichia coli K-12 interacts with the outer membrane peptidoglycan-associated proteins Pal, Lpp and OmpA. Molecular microbiology, 1998. 29(1): p. 359–367.

46. Rassam, P., et al., Supramolecular assemblies underpin turnover of outer membrane proteins in bacteria. Nature, 2015. 523(7560): p. 333–336.

47. Casadaban, M.J., Transposition and fusion of the lac genes to selected promoters in Escherichia coli using bacteriophage lambda and Mu. Journal of molecular biology, 1976. 104(3): p. 541–555.

48. Ruiz, N., et al., Identification of two inner-membrane proteins required for the transport of lipopolysaccharide to the outer membrane of Escherichia coli. Proceedings of the National Academy of Sciences, 2008. 105(14): p. 5537–5542.

49. Baba, T., et al., Construction of Escherichia coli K-12 in-frame, single-gene knockout mutants: the Keio collection. Molecular systems biology, 2006. 2(1): p. 2006.0008.

50. Datsenko, K.A. and B.L. Wanner, One-step inactivation of chromosomal genes in Escherichia coli K-12 using PCR products. Proceedings of the National Academy of Sciences, 2000. 97(12): p. 6640–6645.

51. Silhavy, T.J., M.L. Berman, and L.W. Enquist, Experiments with gene fusions. 1984: Cold Spring Harbor Laboratory.

52. Zomer, A., et al., ESSENTIALS: software for rapid analysis of high throughput transposon insertion sequencing data. 2012.

53. Laehnemann, D., et al., Genomics of rapid adaptation to antibiotics: convergent evolution and scalable sequence amplification. Genome biology and evolution, 2014. 6(6): p. 1287–1301.

