## Supplementary figures and tables for "Genetic interaction mapping highlights key roles of the Tol-Pal complex"

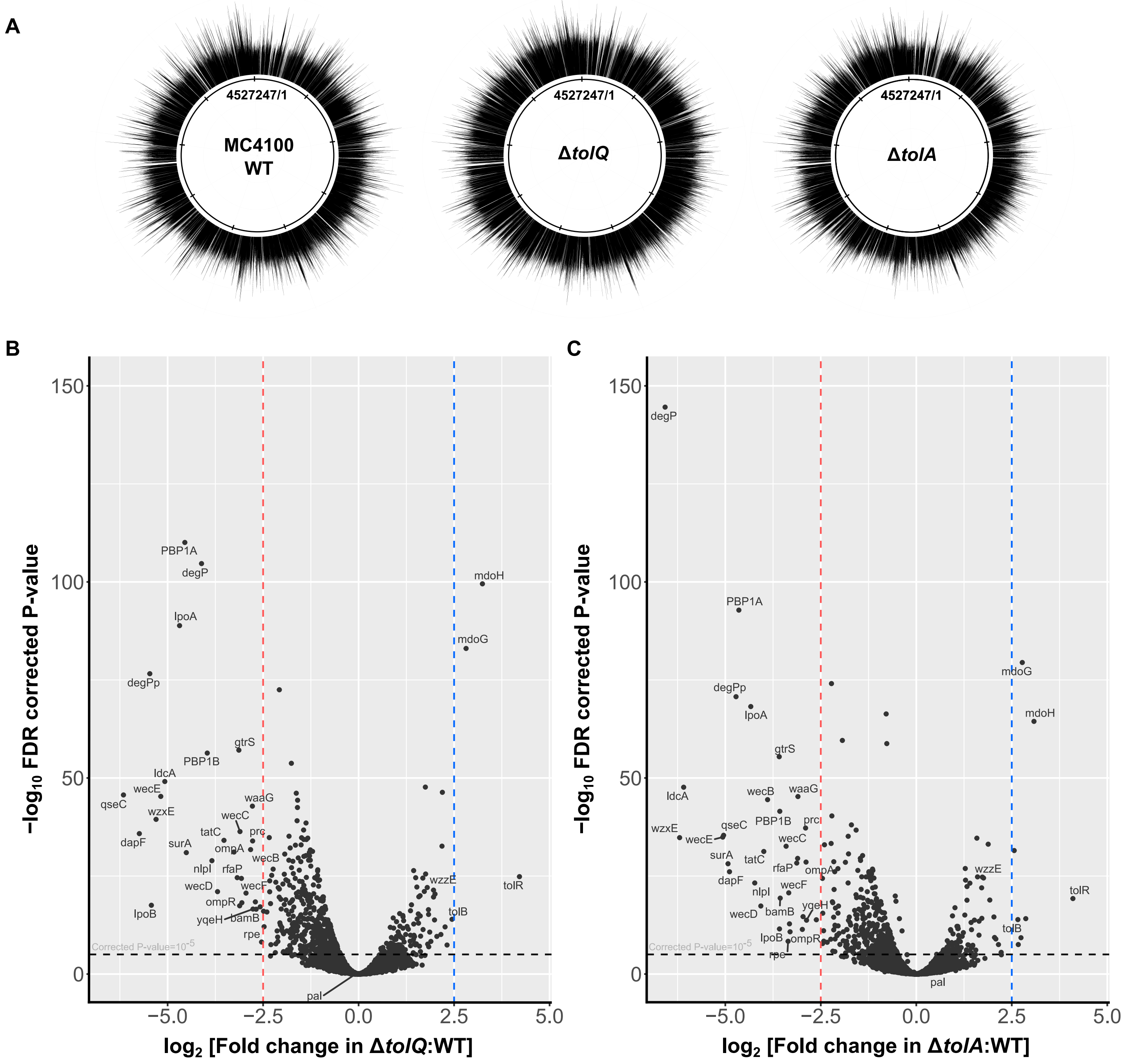


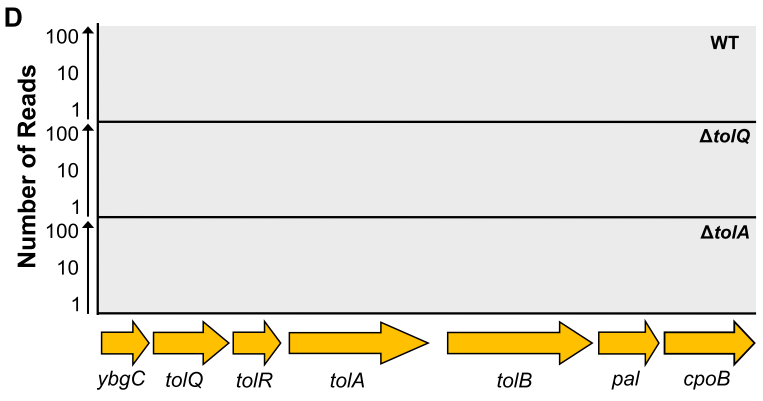


Log$_2$FC of Tn5 insertion frequency at the *tol-pal* locus

| | *ybgC* | *tolQ* | *tolR* | *tolA* | *tolB* | *pal* | *cpoB* |
|---|---|---|---|---|---|---|---|
| *tolQ*:WT | -0.42 | N.A. | **4.21** | **3.12** | **2.45** | -0.07 | -0.21 |
| *tolA*:WT | 0.37 | **2.07** | **4.10** | N.A. | **2.66** | 0.32 | -0.58 |

Greyed out values are not statistically significant

**Figure S1 Saturated transposon libraries allow identification of genetic interactions with the *tol-pal* locus.** (A) WT, Δ*tolQ*, and Δ*tolA* libraries are saturated with transposon insertions randomly distributed across the genome. Frequencies of transposon insertions at each unique site are plotted on the genome map of the respective backgrounds. (B, C) Volcano plots showing statistical significance (corrected P-value) against $\log_2$ fold change ($\log_2$FC) of transposon insertion frequency in each gene for (B) Δ*tolQ* relative to WT and (C) Δ*tolA* relative to WT. Genes with $\log_2$FC more than 2.5 (blue dotted line) or less than -2.5 (red dotted line) are considered strong positive or negative genetic interactions, respectively. Genetic interactions with corrected P-value of less than $10^{-5}$ (above the horizontal black dotted line) are considered statistically significant. (D) Transposon insertion profiles at the *tol-pal* locus in WT, Δ*tolQ* and Δ*tolA* backgrounds. Numbers of reads (raw data before normalization for downstream analysis) on the genome are plotted with open reading frames annotated below the horizontal axis.

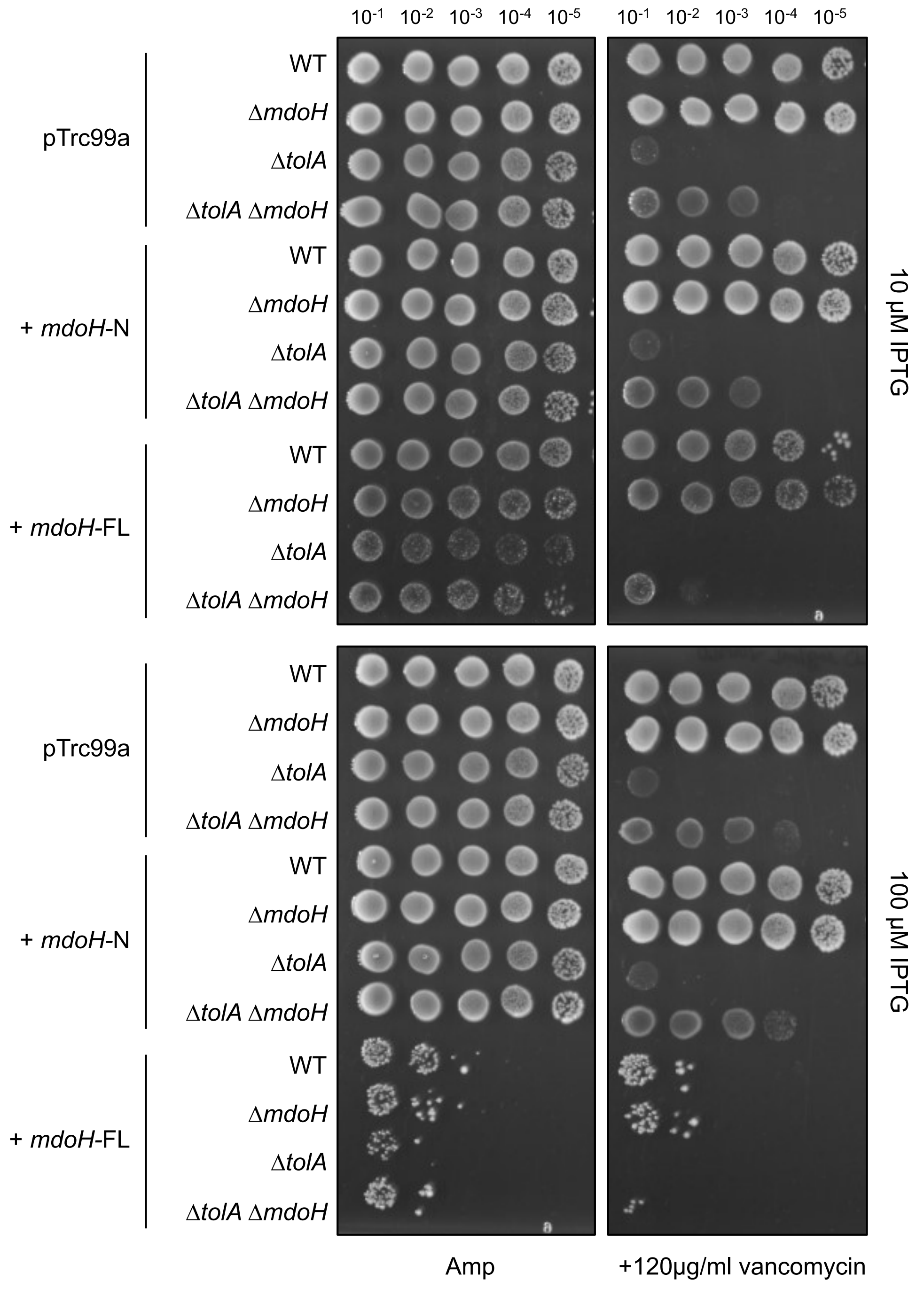


**Figure S2 Overexpressing the N-terminal cytoplasmic domain of MdoH is not sufficient to reverse the suppression of vancomycin sensitivity in the Δ*tolA* strain by Δ*mdoH*.** Efficiency of plating (EOP) of the indicated strains, harbouring either pTrc99a empty vector or pTrc99a-*mdoH*-N/*mdoH*-FL, on LB with vancomycin at 37°C.

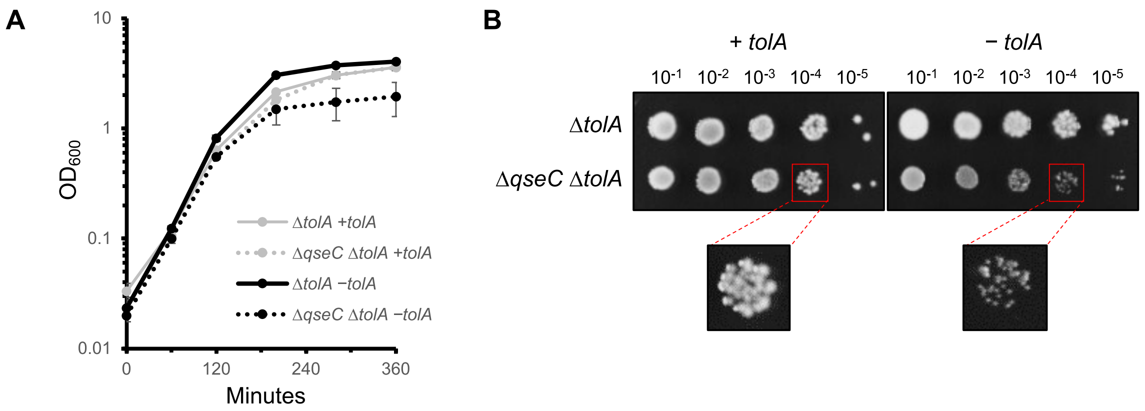


**Figure S3 Depletion of *tolA* results in growth defects in the Δ*qseC* strain.** (A) Growth curves of indicated strains, based on optical density at 600 nm, in LB media at 37°C. Mid-log cultures of NR754 Δ*tolA* and Δ*tolA* Δ*qseC* strains carrying pBAD43-*tolA* were washed three times and used to inoculate fresh cultures (at $OD_{600}$ = 0.02) either with 0.02% (w/v) arabinose (+*tolA*) or 0.2% (w/v) glucose (−*tolA*). (B) EOP of the indicated strains on LB with 0.02% arabinose (+*tolA*) or 0.2% glucose (−*tolA*) at 37°C. The inoculums from (A) were normalized to $OD_{600}$ of 0.1, serially diluted and spotted on to agar plates.

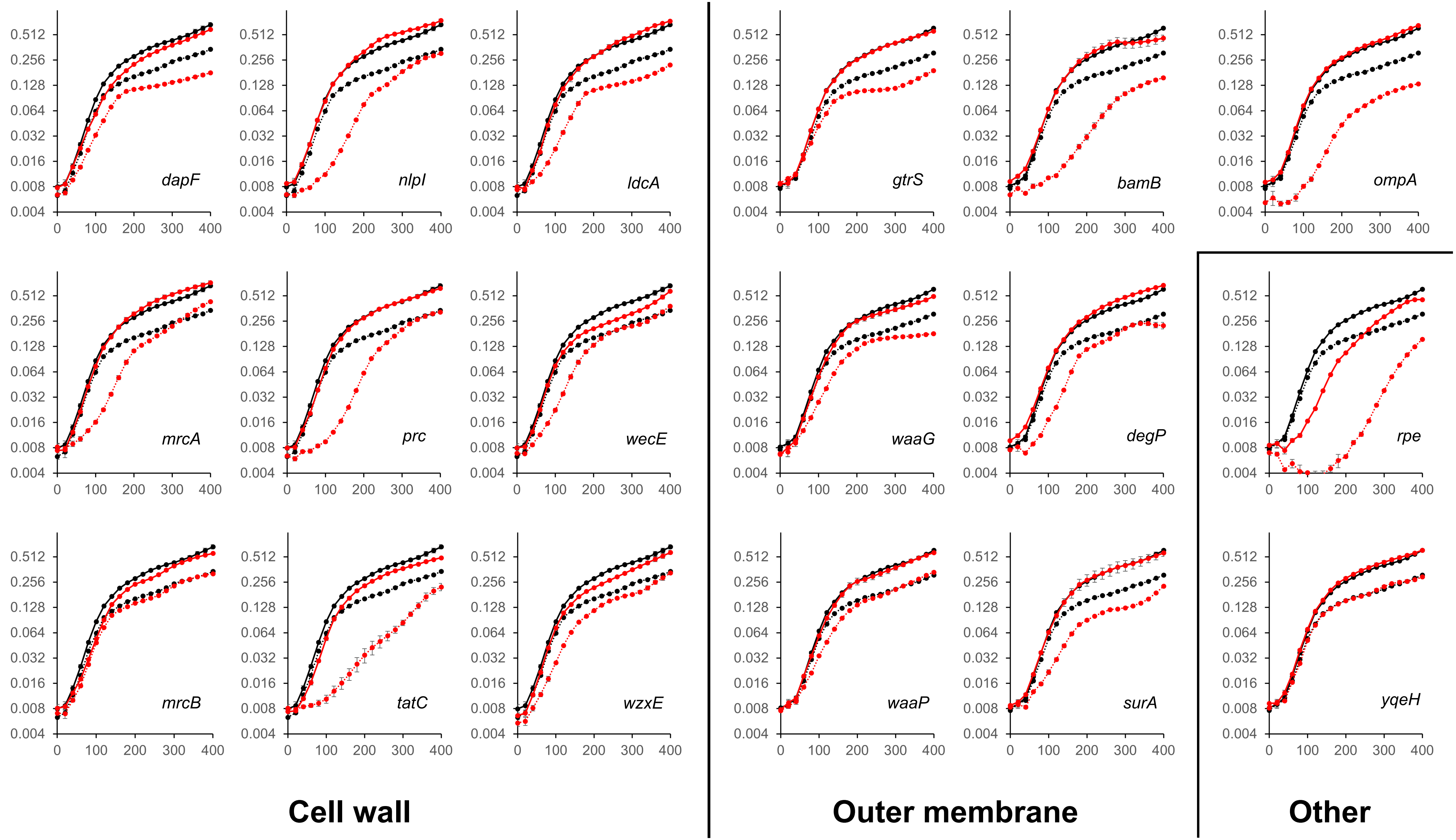


**Figure S4 Negative interactions based on TraDIS correlate with synthetic growth defects in Δ*tolA* background.** Growth curves of indicated strains, based on optical density at 490 nm in 96-wells plates, in LB media at 37°C. WT and Δ*tolA* curves are represented by black solid and dotted lines, while single (in WT background) and double (in Δ*tolA* background) deletion mutants of the indicated gene are represented by red solid and dotted lines, respectively.

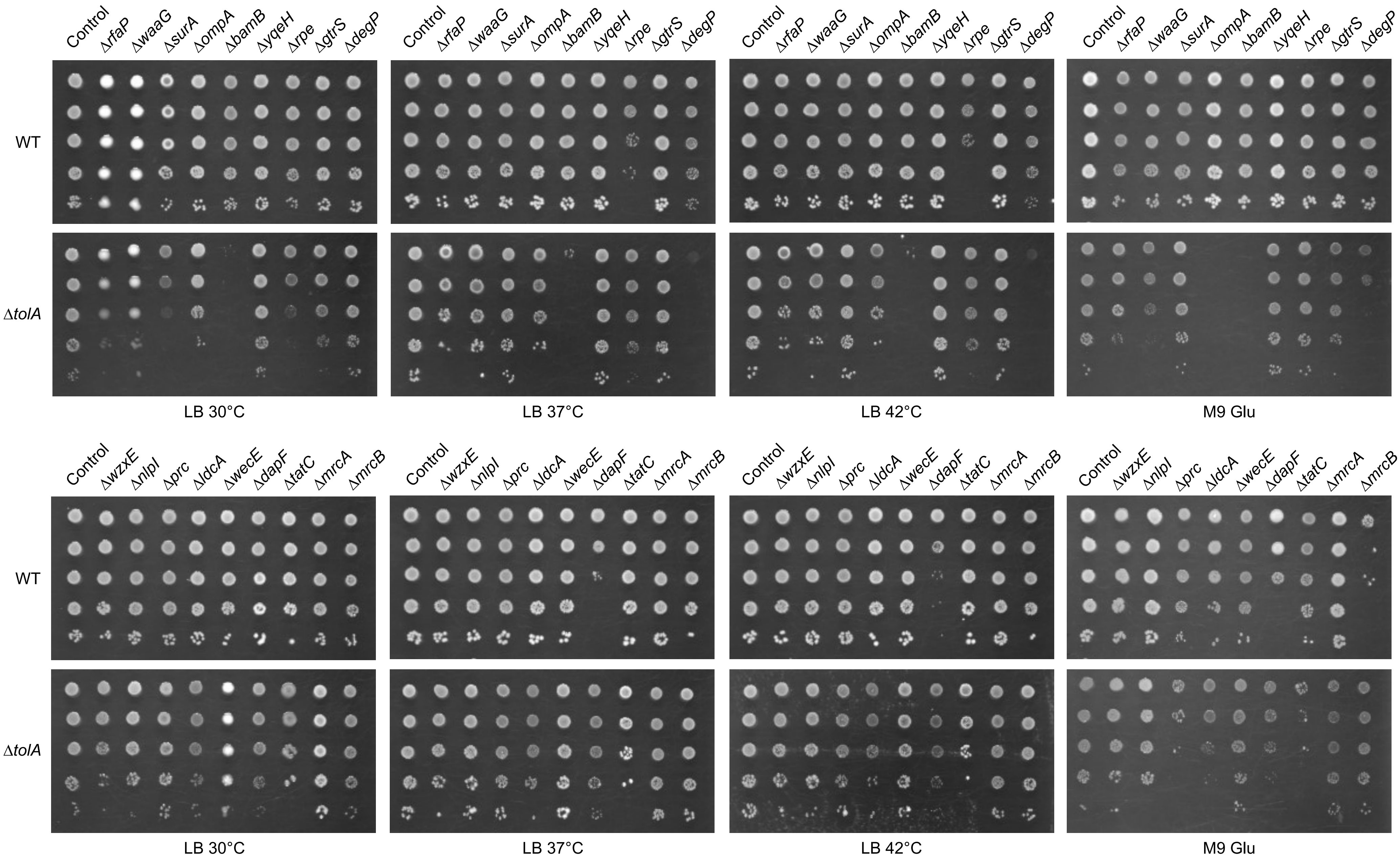


**Figure S5 Negative interactions based on TraDIS correlate with synthetic plating defects in Δ*tolA* background.** EOP of the indicated strains on LB or M9 minimal media at 30°C, 37°C, or 42°C.

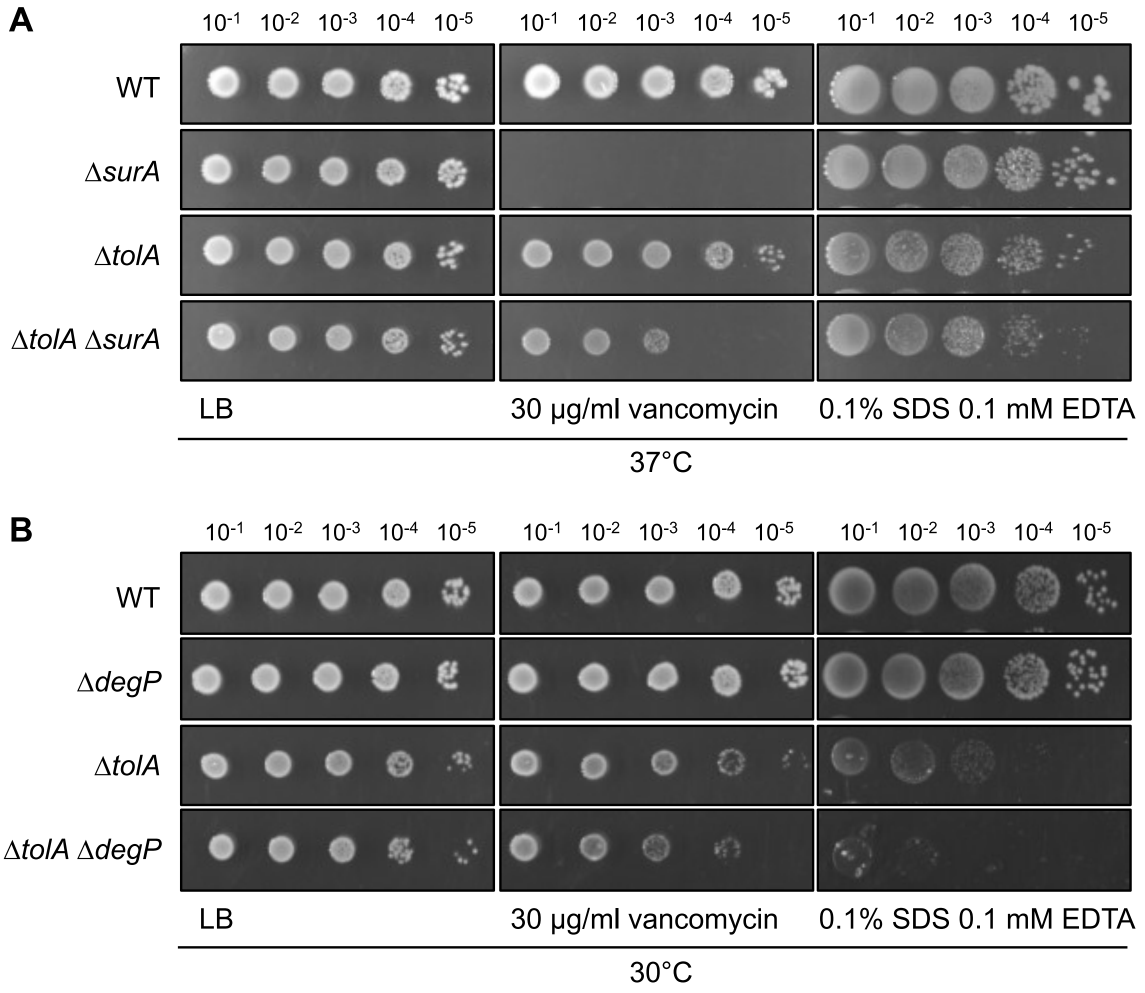


**Figure S6 SDS-EDTA and vancomycin sensitivity of Δ*tolA* Δ*surA* and Δ*tolA* Δ*degP* mutants.** (A) EOP of the indicated strains on LB with vancomycin or SDS-EDTA at 37°C. (B) EOP of the indicated strains on LB with vancomycin or SDS-EDTA at 30°C.

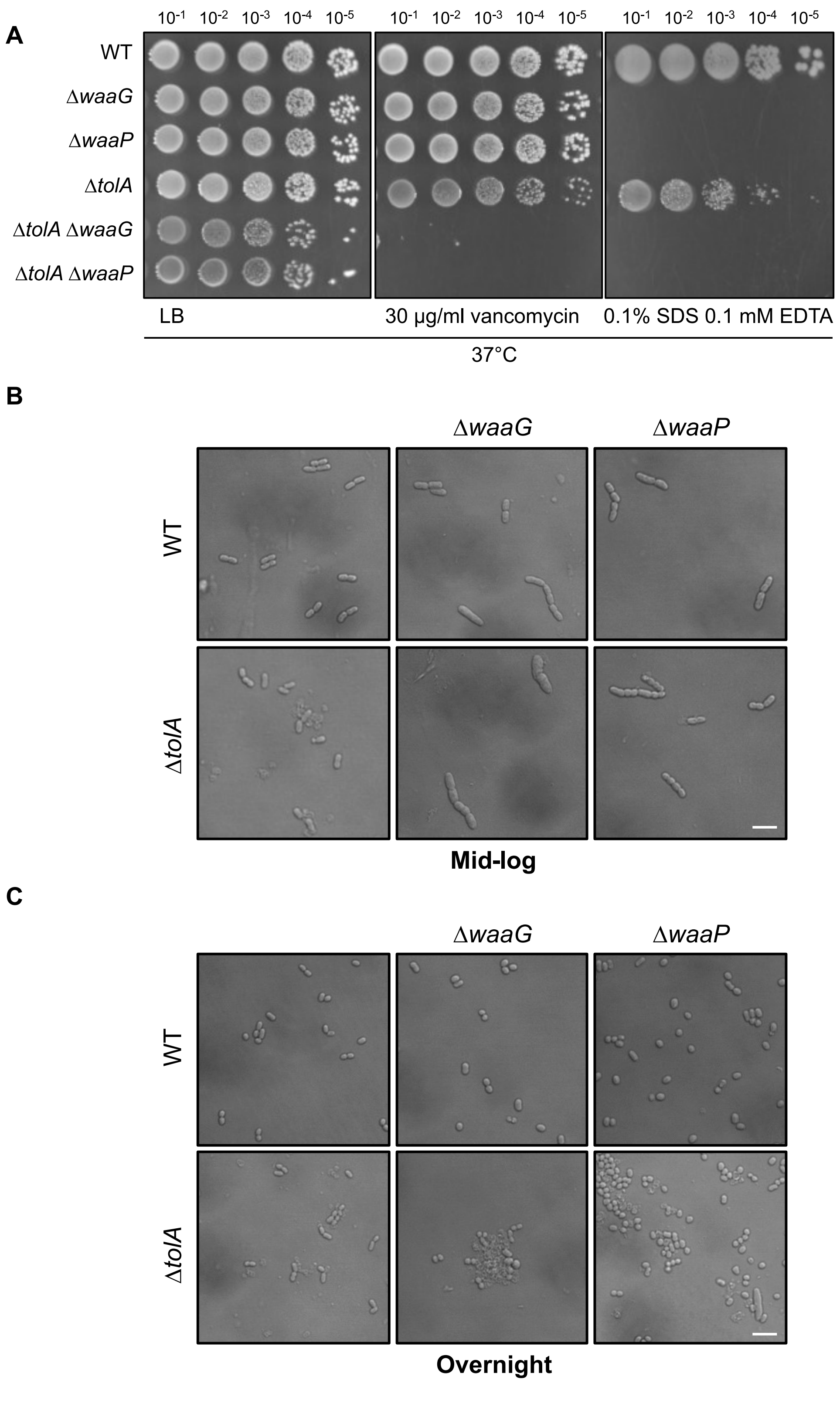


**Figure S7 Defective core LPS biosynthesis is not well-tolerated in Δ*tolA* strains.** (A) EOP of the indicated strains on LB with vancomycin or SDS-EDTA at 37°C. (B, C) DIC images of cells from indicated strains grown to (B) mid-log or (C) stationary phase in LB media (with NaCl) at 37°C. Scale bar represents 5 µm.

**Figure S1 Saturated transposon libraries allow identification of genetic interactions with the *tol-pal* locus.** (A) WT, Δ*tolQ*, and Δ*tolA* libraries are saturated with transposon insertions randomly distributed across the genome. Frequencies of transposon insertions at each unique site are plotted on the genome map of the respective backgrounds. (B, C) Volcano plots showing statistical significance (corrected P-value) against $\log_2$ fold change ($\log_2$FC) of transposon insertion frequency in each gene for (B) Δ*tolQ* relative to WT and (C) Δ*tolA* relative to WT. Genes with $\log_2$FC more than 2.5 (blue dotted line) or less than -2.5 (red dotted line) are considered strong positive or negative genetic interactions, respectively. Genetic interactions with corrected P-value of less than $10^{-5}$ (above the horizontal black dotted line) are considered statistically significant. (D) Transposon insertion profiles at the *tol-pal* locus in WT, Δ*tolQ* and Δ*tolA* backgrounds. Numbers of reads (raw data before normalization for downstream analysis) on the genome are plotted with open reading frames annotated below the horizontal axis.

**Figure S2 Overexpressing the N-terminal cytoplasmic domain of MdoH is not sufficient to reverse the suppression of vancomycin sensitivity in the Δ*tolA* strain by Δ*mdoH*.** Efficiency of plating (EOP) of the indicated strains, harbouring either pTrc99a empty vector or pTrc99a-*mdoH*-N/*mdoH*-FL, on LB with vancomycin at 37°C.

**Figure S3 Depletion of *tolA* results in growth defects in the Δ*qseC* strain.** (A) Growth curves of indicated strains, based on optical density at 600 nm, in LB media at 37°C. Mid-log cultures of NR754 Δ*tolA* and Δ*tolA* Δ*qseC* strains carrying pBAD43-*tolA* were washed three times and used to inoculate fresh cultures (at $OD_{600}$ = 0.02) either with 0.02% (w/v) arabinose (+*tolA*) or 0.2% (w/v) glucose (−*tolA*). (B) EOP of the indicated strains on LB with 0.02% arabinose (+*tolA*) or 0.2% glucose (−*tolA*) at 37°C. The inoculums from (A) were normalized to $OD_{600}$ of 0.1, serially diluted and spotted onto agar plates.

**Figure S4 Negative interactions based on TraDIS correlate with synthetic growth defects in Δ*tolA* background.** Growth curves of indicated strains, based on optical density at 490 nm in 96-wells plates, in LB media at 37°C. WT and Δ*tolA* curves are represented by black solid and dotted lines, while single (in WT background) and double (in Δ*tolA* background) deletion mutants of the indicated gene are represented by red solid and dotted lines, respectively.

**Figure S5 Negative interactions based on TraDIS correlate with synthetic plating defects in Δ*tolA* background.** EOP of the indicated strains on LB or M9 minimal media at 30°C, 37°C, or 42°C.

**Figure S6 SDS-EDTA and vancomycin sensitivity of Δ*tolA* Δ*surA* and Δ*tolA* Δ*degP* mutants.** (A) EOP of the indicated strains on LB with vancomycin or SDS-EDTA at 37°C. (B) EOP of the indicated strains on LB with vancomycin or SDS-EDTA at 30°C.

**Figure S7 Defective core LPS biosynthesis is not well-tolerated in Δ*tolA* strains.** (A) EOP of the indicated strains on LB with vancomycin or SDS-EDTA at 37°C. (B, C) DIC images of cells from indicated strains grown to (B) mid-log or (C) stationary phase in LB media (with NaCl) at 37°C. Scale bar represents 5 µm.

**Table S1 Summary of the sequencing and ESSENTIALS analysis output.** As separate file.

**Table S2 Full list of *tolA* genetic interactions based on differential frequency of transposon insertions in Δ*tolA* relative to WT background.**

**Table S3 Full list of *tolQ* genetic interactions based on differential frequency of transposon insertions in Δ*tolQ* relative to WT background.**

**Table S4 List of positive genetic interactions with $\log_2$FC between 2.5 and 1.25 common between Δ*tolQ* and Δ*tolA* datasets.**

**Table S5 List of strains used in this study**

| Strain and genotype | References |
|---|---|
| MC4100 [*F- araD139 Δ(argF-lac) U169 rpsL150 relA1 flbB5301 ptsF25 deoC1 ptsF25 thi*] | Casadaban, 1976 |
| MC4100 Δ*tolA*::FRT Δ*fhuA*::FRT | This study |
| MC4100 Δ*tolQ*::FRT Δ*fhuA*::FRT | This study |
| MC4100 Δ*mdoH*::$cam^R$ | This study |
| MC4100 Δ*yqeH*::$kan^R$ | This study |
| MC4100 Δ*waaG*::$kan^R$ | This study |
| MC4100 Δ*rfaP*::$kan^R$ | This study |
| MC4100 Δ*gtrS*::$kan^R$ | This study |
| MC4100 Δ*rpe*::$kan^R$ | This study |
| MC4100 Δ*surA*::$kan^R$ | This study |
| MC4100 Δ*ompA*::$kan^R$ | This study |
| MC4100 Δ*bamB*::$kan^R$ | This study |
| MC4100 Δ*degP*::$kan^R$ | This study |
| MC4100 Δ*wzxE*::$kan^R$ | This study |
| MC4100 Δ*wecE*::$cam^R$ | This study |
| MC4100 Δ*mrcB*::$kan^R$ | This study |
| MC4100 Δ*dapF*::$kan^R$ | This study |
| MC4100 Δ*ldcA*::$kan^R$ | This study |
| MC4100 Δ*mrcA*::$kan^R$ | This study |
| MC4100 Δ*tatC*::$kan^R$ | This study |
| MC4100 Δ*nlpI*::$kan^R$ | This study |
| MC4100 Δ*prc*::$kan^R$ | This study |
| MC4100 Δ*tolA*::FRT | This study |
| MC4100 Δ*tolA*::FRT Δ*mdoH*::$cam^R$ | This study |
| MC4100 Δ*tolA*::FRT Δ*yqeH*::$kan^R$ | This study |
| MC4100 Δ*tolA*::FRT Δ*waaG*::$kan^R$ | This study |
| MC4100 Δ*tolA*::FRT Δ*rfaP*::$kan^R$ | This study |
| MC4100 Δ*tolA*::FRT Δ*gtrS*::$kan^R$ | This study |
| MC4100 Δ*tolA*::FRT Δ*rpe*::$kan^R$ | This study |
| MC4100 Δ*tolA*::FRT Δ*surA*::$kan^R$ | This study |
| MC4100 Δ*tolA*::FRT Δ*ompA*::$kan^R$ | This study |
| MC4100 Δ*tolA*::FRT Δ*bamB*::$kan^R$ | This study |
| MC4100 Δ*tolA*::FRT Δ*degP*::$kan^R$ | This study |
| MC4100 Δ*tolA*::FRT Δ*wzxE*::$kan^R$ | This study |
| MC4100 Δ*tolA*::FRT Δ*wecE*::$cam^R$ | This study |
| MC4100 Δ*tolA*::FRT Δ*mrcB*::$kan^R$ | This study |
| MC4100 Δ*tolA*::FRT Δ*dapF*::$kan^R$ | This study |
| MC4100 Δ*tolA*::FRT Δ*ldcA*::$kan^R$ | This study |

| | |
|---|---|
| MC4100 Δ*tolA*::FRT Δ*mrcA*::kan$^R$ | This study |
| MC4100 Δ*tolA*::FRT Δ*tatC*::kan$^R$ | This study |
| MC4100 Δ*tolA*::FRT Δ*nlpI*::kan$^R$ | This study |
| MC4100 Δ*tolA*::FRT Δ*prc*::kan$^R$ | This study |

**Table S6 List of primers used for constructing gene deletions**

| Primer name | Sequence |
|---|---|
| nlpIKO5 | AGGACGTTCATTCAACCGTGGTCTTCGGGAGTGGGAAatgATTCCGGGGATCCGTCGACC |
| nlpIKO3 | GGGCTGATGTGTACGTCAGctaTTGCTGGTCCGATTCTGCTGTAGGCTGGAGCTGCTTCG |
| prcKO5 | GAACACCTGGTGTTCTGAAACGGAGGCCGGGCCAGGCatgATTCCGGGGATCCGTCGACC |
| prcKO3 | TTCTTGTGCCTGATTGATAttaCTTGACGGGAGCGGGTTGTGTAGGCTGGAGCTGCTTCG |
| degPKO5 | GCAATTTTGCGTTATCTGTTAATCGAGACTGAAATACatgATTCCGGGGATCCGTCGACC |
| degPKO3 | AGGAAGGGGTTGAGGGAGAttaCTGCATTAACAGGTAGATTGTAGGCTGGAGCTGCTTCG |
| rfaPKO5 | CCAGAAAAAGCCGCGGATATCATTACAGGTGGTTTAGatgATTCCGGGGATCCGTCGACC |
| rfaPKO3 | TACATACTAATAAATATTTttaTAATCCTTTGCGTTGTGTTGTAGGCTGGAGCTGCTTCG |
| waaGKO5 | CAGAAGATGCCCCTTCAGCTGACAGGAATGCACAATTatgATTCCGGGGATCCGTCGACC |
| waaGKO3 | GGCAAGCGGCTCTTTTAATtcaACCATCTAAACCACCTGTTGTAGGCTGGAGCTGCTTCG |
| yqeHKO5 | AATTATGCAATTATATACGGATAGGGAGGTTCTTAACatgATTCCGGGGATCCGTCGACC |
| yqeHKO3 | TCTGGCTGGTCGTTGAGTAtcaATGTTGGACCGAATGTGATGTAGGCTGGAGCTGCTTCG |
| rpeKO5 | TTTCACCCGCGAAAAAATAATTCTCAAGGAGAAGCGGatgATTCCGGGGATCCGTCGACC |
| rpeKO3 | GCCGCGAATATCTTCAAACttaTTCATGACTTACCTTTGCTGTAGGCTGGAGCTGCTTCG |
| gtrSKO5 | AAAAACGCCCGAAATACATCATCAAGAGAGTCAAAAAatgATTCCGGGGATCCGTCGACC |
| gtrSKO3 | ATTGGCGCGCAATTTAAACttaGTGCTTTACATCGCTATTTGTAGGCTGGAGCTGCTTCG |
| dapFKO5 | AAAGTCAGTTTCTGTACCCGCGTGATTGGAGTAAATGatgATTCCGGGGATCCGTCGACC |
| dapFKO3 | AGTTCTTCCCCTGGTTGCTtcaTAGATGAATAAATCCGTCTGTAGGCTGGAGCTGCTTCG |
| ompAKO5 | TTCATGGCGTATTTTGGATGATAACGAGGCGCAAAAAatgATTCCGGGGATCCGTCGACC |
| ompAKO3 | TTTTCTACCAGACGAGAACttaAGCCTGCGGCTGAGTTACTGTAGGCTGGAGCTGCTTCG |
| bamBKO5 | ATGAAAATTAATAATTTGTCCATCTGAGAGGGACCCGatgATTCCGGGGATCCGTCGACC |
| bamBKO3 | AAGTGAACGACAGAGACGAttaACGTGTAATAGAGTACACTGTAGGCTGGAGCTGCTTCG |
| surAKO5 | CGTAATCCGCAGTGCGGTTAATTGAAATGGAAAAAGTATGATTCCGGGGATCCGTCGACC |
| surAKO3 | ACACGTTGGGTTTTAACCATTAGTTGCTCAGGATTTTAACTGTAGGCTGGAGCTGCTTCG |
| ldcAKO5 | ATATACATTTCGTTTCTGTTGTCAGCAAGGAATTGCCATGATTCCGGGGATCCGTCGACC |
| ldcAKO3 | GCCCTGAAGCGTGATTTTTTTACATTTTAAGAACAGGATGTGTAGGCTGGAGCTGCTTCG |
| mrcBKO5 | ATCGGGCTTTTGCGCCTGAATATTGCGGAGAAAAAGCATGATTCCGGGGATCCGTCGACC |
| mrcBKO3 | GGTATTTCACGCTTAGATGTTAATTACTACCAAACATATCTGTAGGCTGGAGCTGCTTCG |
| mrcAKO5 | ATAAACTGCCCAAATGAAACTAAATGGGAAATTTCCAGTGATTCCGGGGATCCGTCGACC |
| mrcAKO3 | CGCCGAAGCGCCTTTTTAATCAGAACAATTCCTGTGCCTCTGTAGGCTGGAGCTGCTTCG |
| tatCKO5 | CTGCACCTTCCCCTTCGTCGAGTGATAAACCGTAAACATGATTCCGGGGATCCGTCGACC |
| tatCKO3 | CCCTGACGGGCGGTTGAATTTATTCTTCAGTTTTTTCGCTTGTAGGCTGGAGCTGCTTCG |
| wzxEKO5 | ACGGTAATTGCGACTTTGTTGAACTACTTTTCCTGATATGATTCCGGGGATCCGTCGACC |
| wzxEKO3 | AGTACGTGAATCAGTACAGTCATGCCCGCCTACGCCAGAGTGTAGGCTGGAGCTGCTTCG |

**Table S7 List of primers used for TraDIS**

| Primer name | Sequence | Remarks |
|---|---|---|
| TnSeqrev | CAAGCAGAAGACGGCATACGAGAT | Used in both PCR |
| TnSeqfwd1 | ACCTGCAGGCATGCAAGCTTCAGG | Used in 1st round of nested PCR to enrich fragments containing Tn5 junction |
| TnSeqfwdAD005 | AATGATACGGCGACCACCGAGATCTACACTCTTTCCCTACACGAC GCTCTTCCGATCTACAGTGAGCTTCAGGGTTGAGATGTGTA | Used in 2nd round of nested PCR to attach sequencing adaptors and barcode |
| TnSeqfwdAD006 | AATGATACGGCGACCACCGAGATCTACACTCTTTCCCTACACGAC GCTCTTCCGATCTGCCAATAGCTTCAGGGTTGAGATGTGTA | |
| TnSeqfwdAD007 | AATGATACGGCGACCACCGAGATCTACACTCTTTCCCTACACGAC GCTCTTCCGATCTCAGATCAGCTTCAGGGTTGAGATGTGTA | |
| TnSeqfwdAD012 | AATGATACGGCGACCACCGAGATCTACACTCTTTCCCTACACGAC GCTCTTCCGATCTCTTGTAAGCTTCAGGGTTGAGATGTGTA | |
| TnSeqfwdAD013 | AATGATACGGCGACCACCGAGATCTACACTCTTTCCCTACACGAC GCTCTTCCGATCTAGTCAAAGCTTCAGGGTTGAGATGTGTA | |
| TnSeqfwdAD014 | AATGATACGGCGACCACCGAGATCTACACTCTTTCCCTACACGAC GCTCTTCCGATCTAGTTCCAGCTTCAGGGTTGAGATGTGTA | |
| TnSeqfwdAD015 | AATGATACGGCGACCACCGAGATCTACACTCTTTCCCTACACGAC GCTCTTCCGATCTATGTCAAGCTTCAGGGTTGAGATGTGTA | |
| TnSeqfwdAD016 | AATGATACGGCGACCACCGAGATCTACACTCTTTCCCTACACGAC GCTCTTCCGATCTCCGTCCAGCTTCAGGGTTGAGATGTGTA | |
| TnSeqfwdAD019 | AATGATACGGCGACCACCGAGATCTACACTCTTTCCCTACACGAC GCTCTTCCGATCTGTGAAAAGCTTCAGGGTTGAGATGTGTA | |
